# Discovery of new genetic determinants controlling the morphological plasticity in rice root and shoot under phosphate starvation using GWAS

**DOI:** 10.1101/2020.10.31.363556

**Authors:** Nga T P Mai, Chung Duc Mai, Hiep Van Nguyen, Khang Quoc Le, Linh Viet Duong, Huong Thi Mai To

**Author notes:** Correspondence author Email adresse (Huong Thi Mai To).

## Abstract

Phosphorus is an essential nutrient for plants that is often in short supply. In rice (*Oryza sativa* L.), phosphate (Pi) deficiency leads to various physiological disorders that consequently affect plant productivity. In this study, a large-scale phenotyping experiment of a set of 160 Vietnamese rice landraces was performed under greenhouse conditions by employing an alpha lattice design with three replicates to identify quantitative trait loci (QTLs) associated with plant growth inhibition by Pi deficiency. Rice plantlets were grown for six weeks in the PVC sand column (16 cm diameter × 80 cm height) supplied with Pi-deficient (10 µM P) medium or full Pi Yoshida (320 µM P) medium. The effects of Pi deficiency on the number of crown roots, root length, shoot length, root weight, shoot weight and total weight were studied. From 36 significant markers identified by using Genome-wide association study, a total of 21 QTLs associated with plant growth inhibition under Pi starvation conditions were defined. A list of 158 candidate genes co-located with defined QTLs was found. Interestingly, a QTL namely *qRST9*.*14* were detected found common across three weight-traits. The co-located gene *GLYCEROPHOSPHODIESTER PHOSPHODIESTERASE 13* was found potentially involved in Pi transport. Understanding the molecular mechanisms of Pi starvation responses, and identifying potential QTLs responsible for low-Pi stress tolerance will provide valuable information for developing new varieties tolerant to low-Pi conditions.

## 1. Introduction

Rice (*Oryza sativa* L.) is a staple food that is consumed daily as the main carbohydrate source by half of the world’s population. Similar to all other food crops, rice growth and productivity depend greatly on the nutrient availability in and uptake from soil through its root system. Phosphorus (P) is an essential nutrient for plants that is often in short supply (Balemi and Negisho, 2012). Plants require P for the development of many important cellular components, such as nucleic acids, cell membrane, energy-carrying molecules, as well as for various metabolic processes, including photosynthesis and cell respiration (Dong et al., 2019). In rice, phosphate (Pi) deficiency leads to various physiological disorders, such as stunted growth, reduced tillering, slender stems and yield reduction (Dobermann and Fairhurst, 2000). Modern agriculture relies heavily on Pi fertilizers, and it is estimated that approximately 52.9 million tons of Pi fertilizers will be used in agriculture by 2030 (Brears, 2015). Nevertheless, Pi rock is a finite, non-renewable resource, and is predicted to be depleted within 50–200 years under the current level of utilization (Herrera-Estrella and López-Arredondo, 2016). Thus, the development of crop varieties with enhanced P use efficiency (PUE) is one of the critical tasks, which faces researchers seeking to address the problem of Pi starvation (Heuer et al., 2017).

The search for plant genes responsible for tolerance to Pi deficiency has been ongoing for many years (Ni et al., 1998; Wissuwa et al., 1998). Genetic diversity in traits related to low-Pi tolerance provides a potential resource from which plant breeders can develop new varieties that use Pi more effectively (Mehra et al., 2015). Thus, quantitative trait locus (QTL) mapping has been carried out for many low-Pi tolerance-related traits in rice (Chin et al., 2011; Lang and Bui, 2006; Li et al., 2009; Ni et al., 1998; Shimizu et al., 2004; Wissuwa et al., 1998). A QTL named *phosphorus uptake 1* (*Pup1*), which is responsible for rice tolerance to low-Pi conditions, is one of the most promising QTLs for the development of rice cultivars with improved tolerance to Pi starvation (Wissuwa et al., 2002). The *PHOSPHORUS STARVATION TOLERANCE 1* (*OsPSTOL1*) gene, which is located in the *Pup1* region of chromosome (Chr) 12 (Gamuyao et al., 2012), is involved in promoting plant growth and Pi uptake efficiency (Gamuyao et al., 2012). Different classical mapping population have been used to identify QTLs associated with low-Pi tolerance-related traits in rice, such as backcross inbred lines (BILs) (Wissuwa et al., 1998), chromosome segment substitution lines (CSSLs) (Shimizu et al., 2008), crosses between tolerant and sensitive lines, and linkage mapping (Lang and Bui, 2006; Shimizu et al., 2004). One study using BILs showed that a QTL located on Chr 12 and linked to marker C443 has a major effect on Pi intake (Wissuwa et al., 1998). This QTL was found to be responsible for 27.9% of the observed variation in Pi uptake and explained for 26.5% of the variation in dry weight under Pi deficiency (Wissuwa et al., 1998). Another study using CSSLs demonstrated that a rice QTL associated with root elongation under Pi deficiency (*qREP-6*, located on Chr 6) accounted for 54.5% of phenotypic variance under Pi deficiency (Shimizu et al., 2008).

Recent progress in high-throughput sequencing technology has offered a vast opportunity to decipher the genetic variations of individuals within a population (Li et al., 2018). Variations in single nucleotide polymorphisms (SNPs), insertions and deletions can enable plants to adapt to unfavorable environmental conditions (Batley et al., 2003). Genome-wide association study (GWAS) can identify associations between genotype and phenotype, and has been used to analyze the effects of genetic variants in response to Pi starvation in *Arabidopsis thaliana* (El-Soda et al., 2019; Wissuwa et al., 2015; D. Zhang et al., 2014). Association analysis and other systems genomic methods of analyzing root growth rates in 277 *A. thaliana* accessions have led to the discovery of three genes that control variations in root growth in response to low-Pi alone or in combination with other nutrient deficiency conditions (Bouain et al., 2019). Kisko et al. (2018) in their GWAS used 223 *A. thaliana* accessions and demonstrated the key role of *LYSOPHOSPHATIDYLCHOLINE ACYLTRANSFERASE 1* (*LPCAT1*) in shoot Pi accumulation under Zn deficiency by up-regulating the expression of the Pi-transporter gene *PHOSPHATE TRANSPORTER* (*PHT)1;1* (Kisko et al., 2018). Following this strategy, a core panel of 182 rice accessions originating from different regions and ecosystems in Vietnam was collected, yielding 21,623 markers *via* genotyping by sequencing (GBS) (Phung et al., 2014). Recently, this collection has been characterized for rice root-related traits (Phung et al., 2016), panicle-related traits (Ta et al., 2018) and jasmonic acid sensitivity (To et al., 2019) using the GWAS approach. However, the genetic variations responsible for the phenotypic plasticity in rice root and shoot growth under Pi starvation conditions has not been investigated for this panel using GWAS.

In this study, we conducted a GWAS of this Vietnamese rice collection with the aim to identify the genetic diversity, QTLs and genes responsible for morphological plasticity of rice under Pi starvation. As a result, numerous SNPs, QTLs and promising genes associated with rice tolerance to Pi starvation were identified. These results provide valuable information that can be further used for functional genomics-assisted crop improvement.

## 2. Materials and Methods

### 2.1 Plant materials and genotyping data

The rice panel used in this study was composed of 160 rice genotypes, the seeds of which were provided by the Plant Resources Center (Hanoi City, Vietnam). This collection was structured into two major subpanels comprising 94 *indica* and 61 *japonica* cultivars, and one minor subpanel including six admixtures. The list of cultivars and their related information are provided (**Appendix Table A1)**. The GBS data yielded 25,971 SNP markers that were previously identified and are publically available at TropGene Database (Phung et al., 2014).

### 2.2 Plant growth conditions

Each of the 160 accessions was separately germinated in seedling bags for 5 days before being transplanted into sand columns. The phenotyping experiments were carried out in a greenhouse at a temperature of 28°C–30°C, and approximately 70%-80% humidity. The experiment was conducted following a randomized complete block design (RCDB) algorism using IRRISTAT v4.0 with three replicates. Polyvinyl chloride (PVC) columns were filled with fine river sand in drainable black plastic bags (16 cm × 80 cm, diameter × height) and each individual plantlet was grown in separated container (Phung et al., 2016). In each replicate, 160 rice accessions were exposed to the following two Pi application rates: full Pi (320 µM P correspond to 9.92 mg.L^-1^ P) as the control condition and low-Pi (10 µM P correspond to 0.31 mg.L^-1^ P) as the treatment condition. For each genotype, 6 rice plantlets were used in which 3 plants were grown under control condition and 3 plants were grown under low Pi condition. Nutrient levels used for the full Pi condition were as described by Yoshida et al. (Yoshida et al., 1971). Specifically, 50 mg/L NaH_2_PO_4_.H_2_O (320 µM P) was used in full Pi medium; whereas 1.56 mg/L NaH_2_PO_4_.H_2_O (10 µM P) was used in the low Pi medium. The volume of solution using for watering is 2 liters of medium for each plant which corresponds to 19.84 mg P and 0.62 mg P in the full Pi or low Pi condition, respectively. Plants were irrigated with 150 mL the full Pi medium for the control or with low-Pi medium for the treatment every three days during six weeks in order to balance to the amount of evaporated water and the drainage water.

### 2.3 Sample harvesting and phenotyping

After six weeks of growth in sand columns, plants were harvested. The plastic bag containing the roots of each plant was carefully opened, and then the root system of each plant was washed under a faucet with very light water pressure to completely remove the sand. All rice plantlets were phenotyped according to six parameters, three of which shoot length (SHL), root length (RTL), and number of crown root (NCR) were measured immediately after harvesting. The length of the longest leaf was recorded as SHL; the maximum root length as RTL; the number of crown roots were counted manually and recorded as NCR. The samples were then dried in an oven at 70°C for one week to obtain the dry weight. The shoots and leaves, including the stem base, were weighed to obtain shoot weight (SHW); the roots were weighed to obtain root weight (RTW); and the total weight (TTW) was calculated by the sum of SHW and RTW. The phenotypic plasticity index or relative ratios of variation was calculated by the difference between the arithmetic means of the trait under P deficiency and trait under full Pi condition then dividing by means of the trait in full Pi condition on six traits.

### 2.4 Genome-wide association study

To determine the genetic architecture of the Vietnamese rice panel, a principal component analysis (PCA) was run on the Hapmap datasets. The first six PCA axes were retained and used as the structure matrix for the entire panel (Phung et al., 2016). The GWAS analysis was conducted using an mixed linear model (MLM). By incorporating both population stratification and kin relationships among all cultivars, this model effectively eliminated false positive rates. The option parameters “no compression” and “re-evaluation of variance components for each marker” were selected. The association between SNP markers and phenotypic traits was analyzed using Trait Analysis by Association, Evolution and Linkage (TASSEL) v5.2.55. The False Discovery rate (FDR) was analyzed using “qvalue” package in R according to the method in (Phung et al. 2016). The *q*-value corresponds to adjusted *p*-value after FDR analysis was computed to estimate the false discovery rate. However, we applied a less stringent *p*-value of 3.0e-4 as a suggestive threshold to declare that an association was significant to reduce the false negative discovery rate (Phung et al. 2016, To et al., 2019). Q-Q plots were generated to graphically evaluate the number of false positives observed with each model based on deviations from uniform distribution; these were also presented using TASSEL v.5.2.55.

### 2.5 Linkage disequilibrium (LD) heat map and haplotype analysis

The pairwise LD between significant SNPs and their surrounding markers were calculated and visualized according to chromosome location using the LDheatmap R package. The *R*^*2*^ values between pairs of SNP markers were computed; these ranged from 0 to 1. Regions were considered QTLs only in LD blocks with *r*^*2*^ > 0.6. Two main haplotypes were then compared with their respective phenotypes in order to assess the effect of sequence variations on trait phenotype.

### 2.6 Candidate genes

The positions of significantly associated SNPs were clustered into defined QTLs. The positions of the QTLs were then used to screen for candidate genes through the MSU Rice Genome Annotation Project Database Release 7 (http://rice.plantbiology.msu.edu) (Kawahara et al., 2013). Candidates genes co-located with defined QTLs were screened within ± 50kb around the significant markers. Transposons, hypothetical proteins, and expressed proteins were not selected. Only potential genes with annotated functions were used to generate the candidate gene list.

### 2.7 Statistical analysis

All statistical tests (including mean, standard deviation, variance coefficient, ANOVA, and Student t-test used in this study were calculated using R software v3.6.

## 3 Results

### 3.1. Divergent phenotypic responses to Pi deficiency

The variability at the phenotypic level in response to low-Pi conditions were evaluated for six traits as illustrated in the **Figure 1**. Under low-Pi treatment, RTL increased in all genotypes, and the highest and lowest increases were 157% and 2.40%, respectively. Unlike the RTL responses, the five remaining traits were negatively affected by the low-Pi conditions overall. SHL moderately affected from low-Pi levels, with the most affected genotype showing a reduction of 65.80%, compared with 8.90% for the least affected genotype. Nearly all genotypes responded negatively to Pi starvation after the first six weeks, as indicated by a reduced NCR. The most affected genotype showed a 76% reduction in NCR, whereas the least affected genotype showed a reduction of only 5.38%. The weight-related traits were more negatively affected than length traits by the low-Pi treatment. The greatest reduction in RTW was 86.70%; 98.25% for SHW; and 97.70% for TTW. In addition, the phenotypic heterogeneity between the *indica* and *japonica* subgroups was also evaluated and presented in **Figure 2**. Interestingly, genotypes belonging to the *japonica* subgroup were relatively less inhibited by Pi starvation than those belonging to the *indica* subgroup, as indicated by NCR and the root RWT and TTW traits.

**Figure 1.**
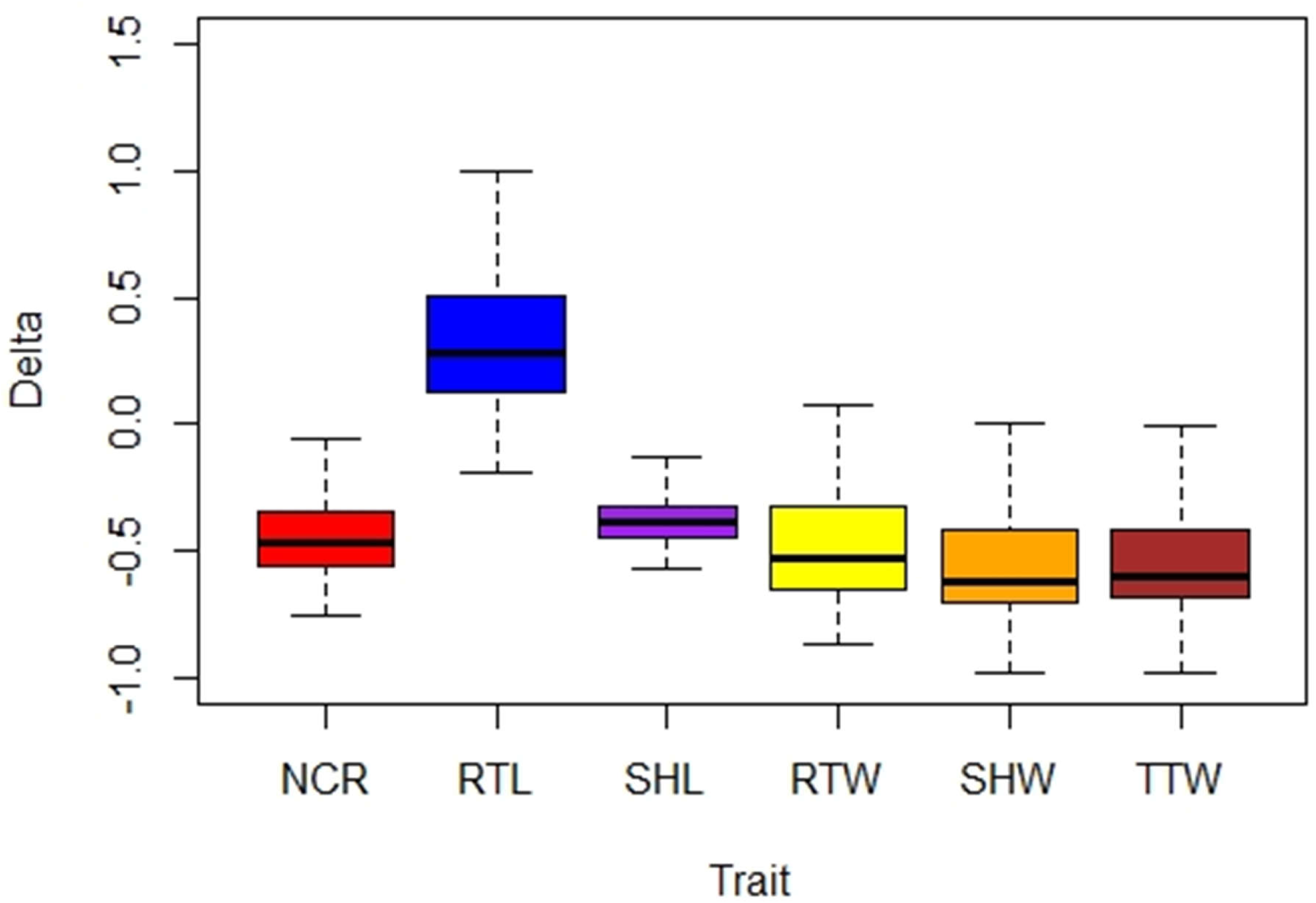
Phenotypic variation in responses to Pi deficiency of six traits. Delta values, or relative ratios of for each of six traits: the number of crown roots (NCR), root length (RTL), shoot length (SHL), root weight (RTW), shoot weight (SHW) and total weight (TTW).

**Figure 2:**
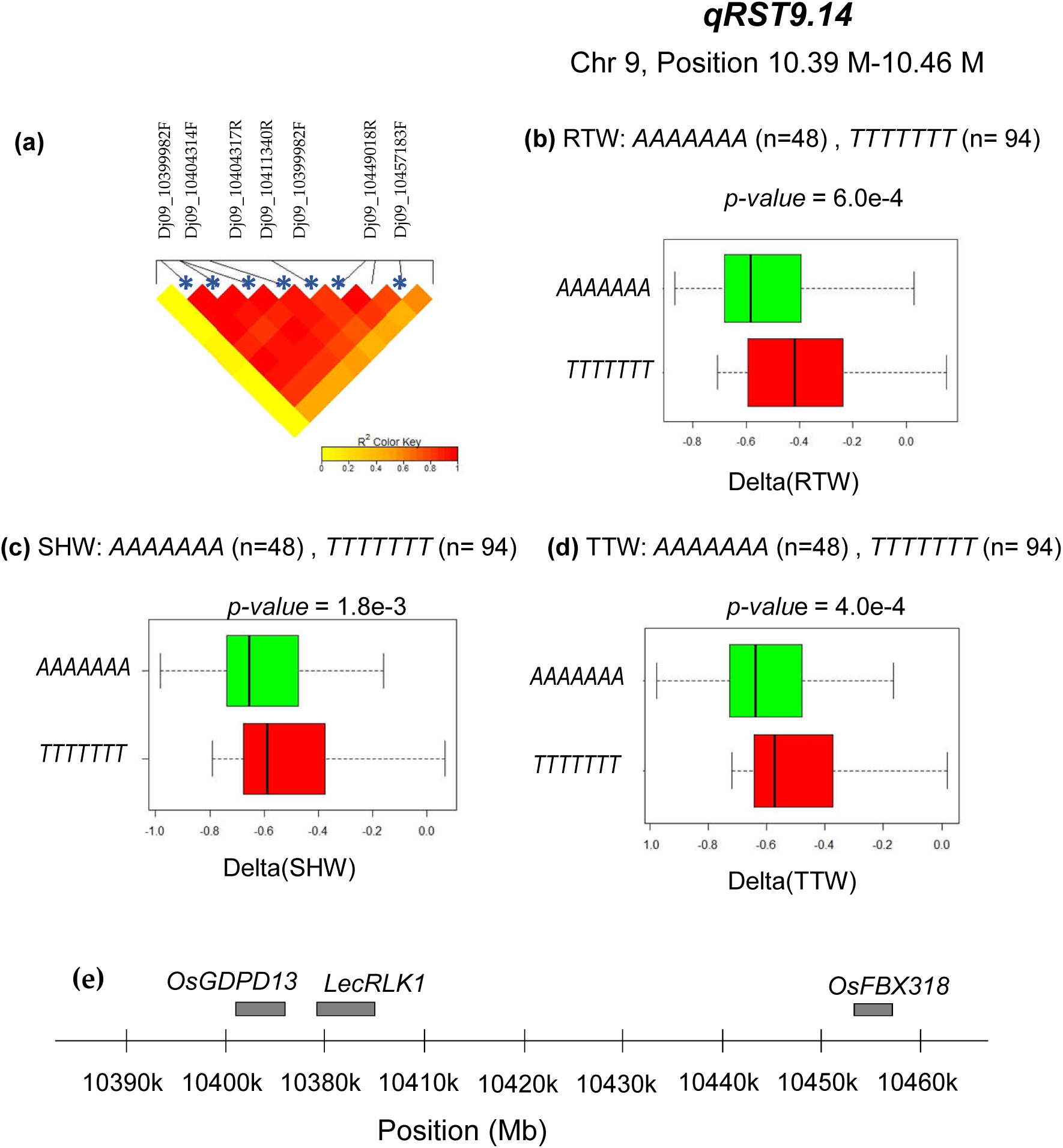
Phenotypic heterogeneity between the *indica* and *japonica* subgroups. Phenotype distribution to (a) number of crown roots (NCR), (b) root length (RTL), (c) root weight (RTW), (d) shoot length (SHL), (e) shoot weight (SHW), and (f) total weight (TTW). Student *t*-test was used to assess the statistically significant differences between each pair of subpanels. Asterisks (*), (**), and (***) correspond to *p* < 0.05, 0.01, and 0.001, respectively.

### 3.2 Association mapping of growth traits in response to Pi starvation and identification of significant markers

The association study was conducted to the 160 Vietnamese rice accessions to determine whether any genetic variants were specifically related to the Pi deficiency response using TASSEL. The MLM, which considers both the kinship matrix and first six principal components (PC) of genotype data, was applied (Phung et al., 2016). Association results obtained using Manhattan plot and Q-Q plot for all six traits are illustrated in **Figure 3**. Overall, the Q-Q plot results concerning the six traits have shown the fitted models between experimental results and theoretical values. The fluctuation tails in the Q-Q plots relate to highly significant *p*-values.

**Figure 3.**
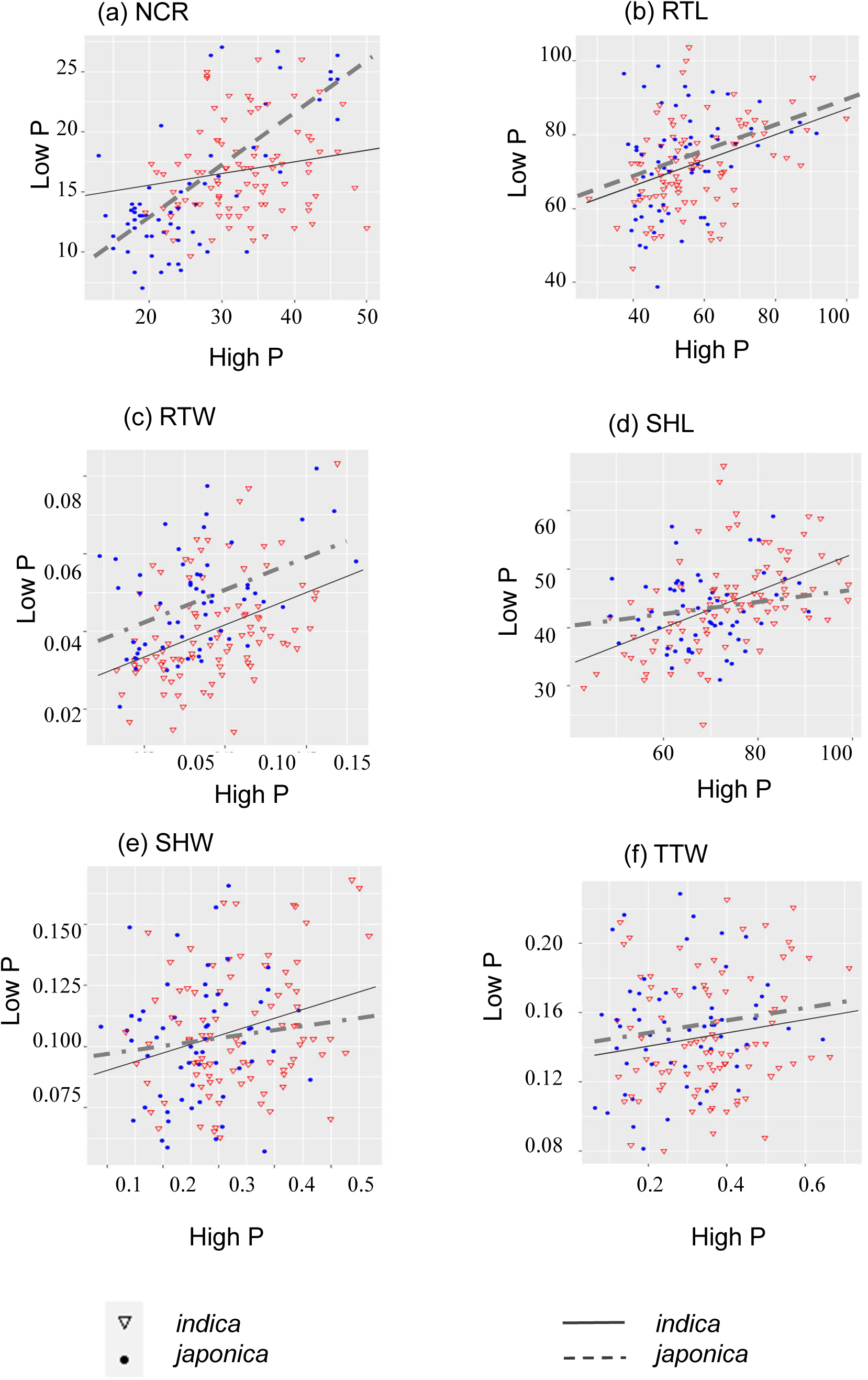
GWAS analysis of the effect of Pi starvation on six traits. Manhattan plots present the distribution of –log (*p*) of 21,623 SNPs markers over 12 chromosomes for (a) shoot weight (SHW), (b) root weight (RTW), (c) total weight (TTW), (d) shoot length (SHL), (e) root length (RTL), and (f) number of crown roots (NCR). The solid line indicates the suggestive significance threshold, *p* = 3.0e-04. The dashed line indicates the threshold with q-values < 0.05. The dot lines indicate the common QTLs across traits.

A total of 36 significant markers (*p* < 3.0e-4) were found to be distributed on eight chromosomes. The number of markers relevant to the NCR, RTL, SHL, RWT, SHW, and TTW traits was 5, 3, 5, 13, 13, and 9, respectively. Detailed information about the positions and *p*-values of the significant markers can be found in **Appendix Table A2**. The statistical results of MLM also revealed some common significant markers among the three weight-related traits (SHW, RTW, and TTW); these share seven common markers located on Chr 9, and one common marker located on Chr 10. Statistical analysis indicated that two most significant association among all markers was Dj09_10399982F and Dj09_10404317R on Chr 9 (*p* = 3.18e-10 for both), which are common across three weight-related traits.

### 3.3 QTL identification

Using the significant SNPs located in each QTL detected using GWAS to perform the LD calculation, the heat maps visually confirmed the strong correlation between each significant SNP and nearby markers. Out of the 36 significant markers identified using GWAS analysis, 21 QTLs related to Pi starvation were identified (**Appendix Table A2)**. Overall, Chr 1 contained the most QTLs, with five QTLs namely *qSHL1*.*1, qSHL1*.*2, qSHW1*.*3, qSHW1*.*4* and *qRTW1*.*5*. The largest QTL (over 300 kb), *qRTL5*.*9*, is located on Chr 5. The smallest (70 kb) is *qSHW9*.*14*, located on Chr 9; this QTL contained seven of the most significant markers.

In total, the NCR trait was associated with five QTLs; the RTL trait with three; the SHL trait with two; SHW and RTW traits with six each. The three weight-related traits shared two common QTLs, namely *qRST9*.*14* (stands for *qRTW9*.*14, qSHW9*.*14* and *qTTW9*.*14*) and *qRST10*.*17* (stands for *qRTW10*.*17, qSHW10*.*17* and *qTTW10*.*17)*. The QTL *qRST9*.*14* was linked to seven common significant markers, namely Dj09_10399982F, Dj09_10404314F, Dj09_10404317R, Dj09_10411340R, Dj09_10426324R, Dj09_10449018R and Dj09_10457183F. The QTL *qRST10*.*17* was linked to one common significant marker including Dj10_21135126R.

### 3.4 Polymorphism combination analysis of significant markers in selected QTLs

The significant SNPs associated with each QTL detected using GWAS were used to assemble a haplotype sequence. In this section, the polymorphism combination analysis of *qRST9*.*14*, which is the most interesting QTLs and appeared common to the TTW, RTW, and SHW traits, was presented in **Figure 4**. The QTL *qRST9*.*14* has seven strongly linked markers as illustrated in **Figure 4a**. The two main haplotypes an *AAAAAAA* and *TTTTTTT* were used to compare the effect of haplotype on a given trait. We found that the accessions containing the *TTTTTTT* sequence variants were significantly less affected by Pi starvation than those containing *AAAAAAA* sequence variants for the RTW trait (**Figure 4b**), SHW trait (**Figure 4c**), and TTW trait (**Figure 4d**). The haplotype analysis was conducted for all 21 QTLs found in this study. Interestingly, the haplotype sequence of 14/21 QTLs was found significantly correlated with their respective phenotypes (**Appendix Figure A1)**.

**Figure 4.**
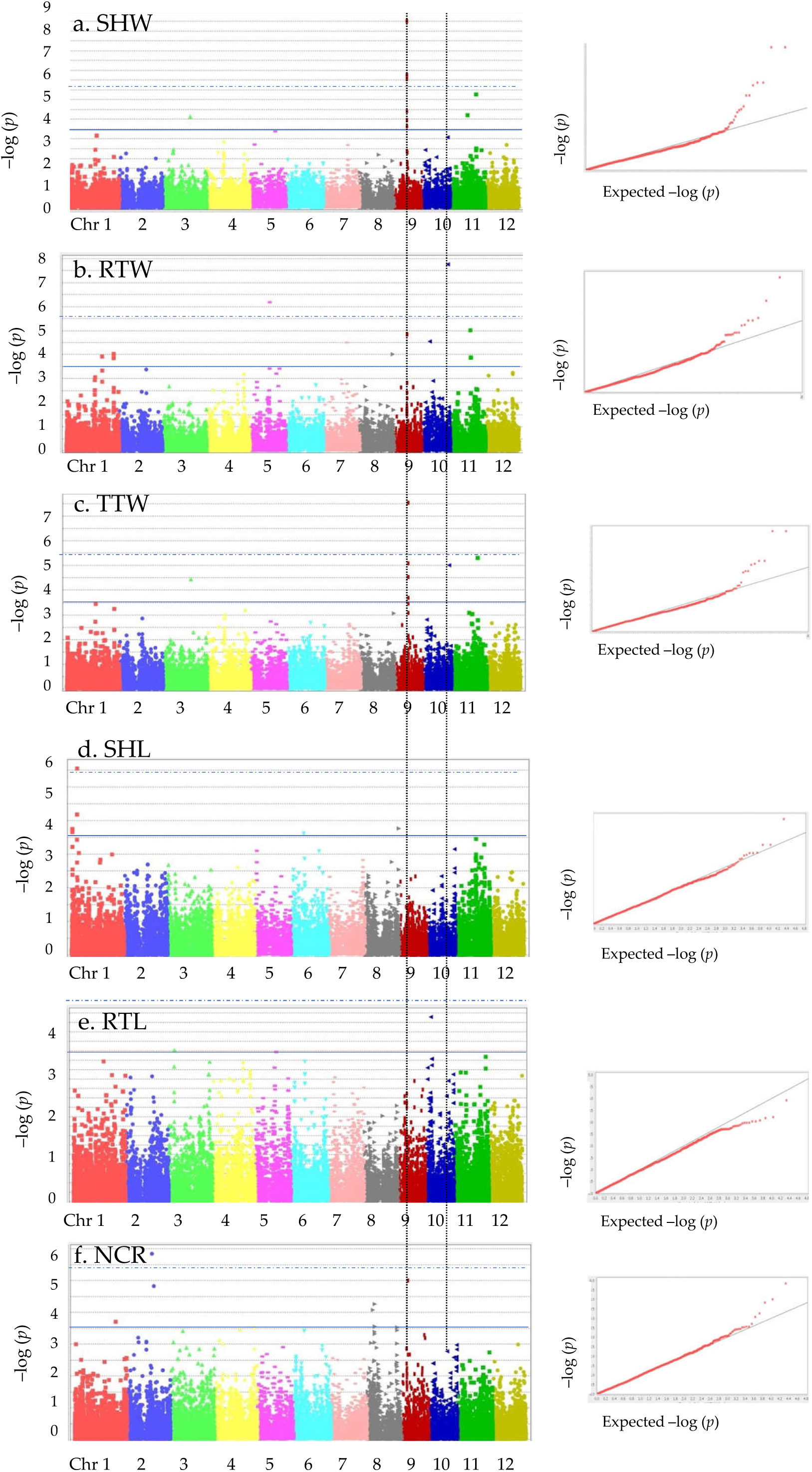
Haplotype analysis for *qRST9*.*14*. (a) Linkage disequilibrium (LD) heat map of the peak region of association for *qRST9*.*14*. Significant SNPs are indicated by a star and the pairwise R^2^ pattern of the associated SNPs in the QTL is indicated with a gradient scale. (b) Effect of allelic combination of two main haplotypes of *qRST9*.*14* on root weight (RTW). (c) Effect of allelic combination of two main haplotypes of *qRST9*.*14* on shoot weight (SHW). (d) Effect of allelic combination of two main haplotypes of *qRST9*.*14* on total weight (TTW). The number of cultivars (n) containing haplotypes *AAAAAAA* and *TTTTTT* is shown. Student’s *t-*test was used to determine significant differences between the two haplotypes. (e) Candidate genes located within QTL interval.

### 3.5 Identification of candidate genes co-localized with significant markers

We looked up the annotations for all candidate genes corresponding to each identified QTL of each trait using the Rice Genome Browser platform of Michigan State University (MSU) Rice Genome Annotation Project Database Release 7. Genes within an interval of 50 kb upstream and downstream of the most significant SNPs derived from GWAS results were examined. From the 21 QTLs identified from 36 significant markers, a total of 158 candidate genes were found (**Appendix Table A2)**. Half of the genes are distributed on Chr 1 and Chr 11. No gene was found on Chr 4, 6, 7, or 12.

From this list, *OsPT1* (LOC_Os01g02000.1) was found to be linked with Pi transport. Many genes were found to be linked with Pi relocalization in the cell, such as *PHOSPHATASES* (LOC_Os01g66920.1, LOC_Os01g08780.1, and LOC_Os10g39540.1) and *GLYCEROPHOSPHORYLDIESTER PHOSPHODIESTERASES* (LOC_Os03g40670 and LOC_Os09g17000).

## 4. Discussion

Although many studies concerning phosphate deficiency tolerance in rice have been conducted, to the best of our knowledge, very few studies have used a panel of GBS rice cultivars to investigate the correlation between natural genetic variation and QTLs related to low-Pi tolerance. In this study, the GWAS approach was applied to identify QTLs associated with plant growth regulation under Pi starvation conditions. We identified 21 significant potential QTLs, including five for NCR, three for RTL, two for SHL, six each for SHW and RTW, and two for TTW. Interestingly, the three weight-related traits shared two common QTLs: *qRST9*.*14* and *qRST10*.*17*. The former harbors seven common significant markers located on Chr 9, and the latter contains one common significant marker located on Chr 10 (**Appendix Table A2**).

In the current GWAS analysis, we identified certain well-known QTLs, which have already been reported in previous studies that used traditional approaches. For example, the QTL *qNCR8*.*13* identified in this study coincides with three QTLs (*qRS8b* [root-shoot ratio], *qRRW8* [relative root dry weight], and *qRRS8* [relative root-shoot ratio]) identified by Li et al. (2009) who used introgression lines (Li et al., 2009). Specifically, three significant markers associated with *qNCR8*.*13* (Sj08_24620154R, Sj08_24632283R, and Dj08_24667998R) are close to the RM5485 and RM6966 markers, which are located within the *qRS8b, qRRW8*, and *qRRS8* regions (Li et al., 2009). The QTL named *qSHW10*.*17* in this study coincides with *qRPH6-qRPH10* (relative plant height) and *qRTDW1-qRTDW10* (relative total dry weight) described by Li et al. (Li et al., 2009). The marker Dj10_21135126R, which we found to be linked to *qSHW10*.*17* in this study, is close to RM5352 and RM333, which are located in *qRPH6-qRPH10* and *qRTDW1-qRTDW10* (Li et al., 2009). Based on this finding, two candidate genes (LOC_Os08g39100.1 and LOC_Os10g39540.1) encoding PHOSPHATASE 2C were found within the *qNCR8*.*13* and *qSHW10*.*17* regions, respectively. These could potentially be the key elements responsible for the functionality of these two confirmed QTLs under low-Pi conditions.

This GWAS was also able to identify some distinct genes already implicated in the Pi starvation regulation pathway in rice. For example, *OsPHO1*.*1* (LOC_Os01g02000.1), which is generally known to be associated with Pi transport (Secco et al., 2010; Wu et al., 2013; Saenchai et al., 2016), was found to co-locate with *qSHL1*.*1*. Five genes encoding for phosphatases (LOC_Os01g08780.1, LOC_Os01g66920.1, LOC_Os03g40670.1, LOC_Os08g39100.1, and LOC_Os10g39540.1), which are related to Pi remobilization (López-Arredondo et al., 2014), were found to co-locate with *qSHL1*.*2, qRTW1*.*4, qSHW3*.*8, qNCR8*.*13*, and *qRST10*.*17*, respectively. The *PHOSPHATE TRANSPORTER 1* (*PHO1*) gene, which was first described in *A. thaliana*, encodes a protein that is known for playing an essential role in transferring Pi from the roots to shoots (Poirier et al., 1991; Secco et al., 2010; Wu et al., 2013). In rice, there are three orthologs to *A. thaliana AtPHO1* which contain *cis*-natural antisense transcripts (Secco et al., 2010), namely *OsPHO1;1, OsPHO1;2* and *OsPHO1;3*. We found *OsPHO1;1* (LOC_Os01g02000.1) to be co-located with *qSHL1*.*1*. Both sense and antisense transcripts of *OsPHO1;2* have been reported to be involved in transporting Pi from roots to shoots in rice (Secco et al., 2010). Moreover, the *cis*-natural antisense transcript of *OsPHO1;1* in rice was up-regulated five-fold under combined Pi–Zn starvation, but not under either stressor individually (Saenchai et al., 2016). In our study, *OsPHO1;1* was detected in the *qSHL1*.*1* region, providing further evidence supporting the involvement of *OsPHO1;1* in the Pi starvation response in rice. Using Genevestigator ® –a curated transcriptomic data from public repositories, we screened all the genes colocalized with the significant markers and found 28/158 of the candidate genes was strongly regulated under low Pi treatment (Appendix Table A3). These results indicated that these genes are potentially important factors in Pi starvation tolerance response in rice. Taken together, the above results firmly support the solidity of the current GWAS analysis. In addition to well-described QTLs and genes, we discovered other significant QTLs and promising genes that have not yet been associated with known functions, but would be very interesting candidates for further investigation.

Transcription factor WRKY30, encoded by *OsWKKY30* (LOC_Os08g38990), is one of ten TFs identified by our GWAS analysis. We found that this gene is located within the *qNCR8*.*13* interval; interestingly, this QTL coincides with three confirmed QTLs mentioned earlier. According to the literature, the *OsWKKY30* gene is involved in both abiotic and biotic stress responses (Peng et al., 2012; Shen et al., 2012). Although no information has been reported regarding the function of *OsWRKY30* in the Pi starvation response, many other genes in the *OsWRKY* and *AtWRKY* families are reported to be involved (Dai et al., 2016; Liu et al., 2011; Wang et al., 2018). The overexpression of *OsWRKY74* in rice significantly enhances the tolerance to Pi starvation, Fe deficiency, and cold stress (Dai et al., 2016). The *OsWRKY74-*overexpressing lines showed increases in root number, shoot biomass, and Pi concentration of approximate 16%; and increases in both tiller number and grain weight of approximate 24% compared with the wild-type (Dai et al., 2016). A lower approximate Pi, lower yield, and altered root systems were obtained in the *oswrky28* mutants compared with wild-type in rice (Wang et al., 2018). In *A. thaliana*, the expression level of *AtWRKY45* increased by up to 22 times more in the roots after five days of treatment with low Pi compared with that under the control condition, and led to an increase in Pi uptake and therefore a higher Pi accumulation in the whole plant (Wang et al., 2014). Further, the RNAi-silencing transgenic lines of *AtWRKY75* accumulated anthocyanin and decreased the expression levels of some Pi–related genes, such as *PHOSPHATASES* (*AtPS2-1, AtPS2-2*) and *PTs* (*AtPht1;1* and *AtPht1;4*) (Devaiah et al., 2007; Z. Zhang et al., 2014). Consequently, we propose that *OsWRKY30* could be a potentially key element in the functionality of *qNCR8*.*13* in Pi-deprived rice.

The GLYCEROPHOSPHODIESTER PHOSPHODIESTERASE (GDPD) enzymes are involved in the metabolic degradation of glycerophosphodiester into sn-glycerol-3-phosphate, which aids in remobilizing glycerophosphate in cells, thereby maintaining phosphate homeostasis (Corda et al., 2014). Numerous GDPD genes are reported to be highly upregulated under Pi starvation conditions in various species, such as in *Arabidopsis*, maize, tomato, and white lupin (Corda et al., 2014; Mehra et al., 2019). In rice, 13 *OsDGPD* genes were identified (Mehra and Giri, 2016). Mehra et al. (2016) characterized the function of this gene family in low-Pi and other nutrient deficiency conditions (e.g., N, K, Fe, and Zn) (Mehra and Giri, 2016). The expression profiles of 12 of the 13 *OsDGPD* genes have been quantified after 7 and 15 days of nutrient starvation, revealing that 11 of these genes are transcriptionally regulated under starvation, particularly by low-Pi conditions. More recent studies have indicated that *OsGDPD2* is transcriptionally regulated by *PHOSPHATE STARVATION RESPONSE (PHR)2* (Mehra et al., 2019). In our GWAS analysis, two *DGPD* genes were detected within the *qSHW3*.*8* and *qRST9*.*14* intervals: *OsDGPD5* (LOC_Os03g40670) and *OsGDPD13* (LOC_Os09g17000), respectively. Mehra et al. (2019) found that the *OsDGPD5* gene responds early to low-Pi treatment (Mehra et al., 2019), and was induced up to 67-fold after 7 days and up to 141-fold after 15 days of treatment. Moreover, *OsDGPD5* is also significantly upregulated after 7 days under low-K (8-fold), low-Fe (4-fold), and low-Zn (1.5-fold) conditions (Mehra and Giri, 2016). This gene was highly upregulated in root tissue under low-Pi treatment compared with full Pi, and might correlate with the genetic background. Specifically, the expression level of *OsDGPD5* was upregulated by as much as 403-fold in *O. sativa* Dular, a tolerant variety, but only 80-fold in *O. sativa* PB1, a sensitive variety (Mehra et al., 2016). Thus, the results obtained by GWAS analysis strongly confirm the important roles of *OsDGPD5* and *qSHW3*.*8* in the low-Pi response. In contrast to the early responsiveness found in *OsGDPD* group A to low-Pi treatment, *OsGDPDs* group B genes were generally late in responding to low-Pi treatment (Cheng et al., 2011). Because the full-length cDNA sequence of *OsGDPD13* was absent from the KOME database, the expression profile of this gene was unavailable in the study of Mehra (Mehra and Giri, 2016). However, it is particularly interesting to note that, according to GWAS analysis, the marker (Dj09_10404317R) located inside the *OsGDPD13* gene had the most significant *p*-value (p = 3.18e-10). The LD heat map analysis showed an extremely strong red block that linked all seven lead markers in this region (**Figure 4a**). Moreover, the significant correlation between haplotype sequence and the corresponding phenotype was found for all SHW, RTW, and TTW traits (**Figure 4b, c and d**). Therefore, *OsGDPD13* and *qRST9*.*14* are very promising targets, thus, need a further in-depth functional characterization.

## 5. Conclusions

This study marks the first time that the Pi deficiency response in rice has been characterized using a thoroughly genotyped Vietnamese rice panel with the GWAS approach. The results reveals some new and robust QTLs (e.g., *qSHW3*.*8, qNCR8*.*13* and *qRST9*.*14*) and some co-located genes (e.g., *OsGPDP5, OsWRKY30* and *OsGPDP13)* within these QTLs, which have a strong potential to regulate the response of rice to Pi starvation conditions. Further investigation of these QTLs and underlying genes by bi-parental mapping and functional genomic analysis will provide more profound understanding of the molecular mechanisms of the Pi deficiency response, and facilitate the breeding and improvement of new rice varieties.

## Supplementary Materials

Supplementary materials can be found at …

## Author Contributions

HTMT and NTPM conceived and designed the experiments. NTPM, DCM, VHN and VLD carried out most of the experiments. HTMT, NTPM and LQK performed bioinformatics analysis. HTMT and NTPM wrote the paper. All authors read and approved the manuscript for publication.

## Funding

This research was funded by Vietnam National Foundation for Science and Technology Development (NAFOSTED) under grant number 106-NN.03-2016.15 to Huong TM To.

## Acknowledgements

We would like give a special thanks to “Rice Functional Genomics and Plant Biology” International Joint Laboratory (LMI-RICE2) and Prof. Michel Lebrun for the financial support for the multiplication of the rice collection. We also extend our deep gratitude to Dr. Lam-Son Phan Tran for his insightful comment for this manuscript. We thank Dr Phung Thi Phuong Nhung for her advice in the phenotyping design and Prof. Luu Thi Cuc (AGI) for the green house. We acknowledge our beloved USTH and VNUA students, including Quynh Anh, Thanh Tung, Robin Jacquemin, Thuy Linh, Ngoc Tit, Le Na, Van Anh, and Phuong Trang, and Manh Tung, for their kind participation in the phenotyping experiments.

## Conflicts of Interest

The authors declare no conflict of interest.

## Notes

### Competing Interest Statement

The authors have declared no competing interest.

