## Supplementary material for "Discovery of new genetic determinants controlling the morphological plasticity in rice root and shoot under phosphate starvation using GWAS": Sup Fig A1

### Slide 1
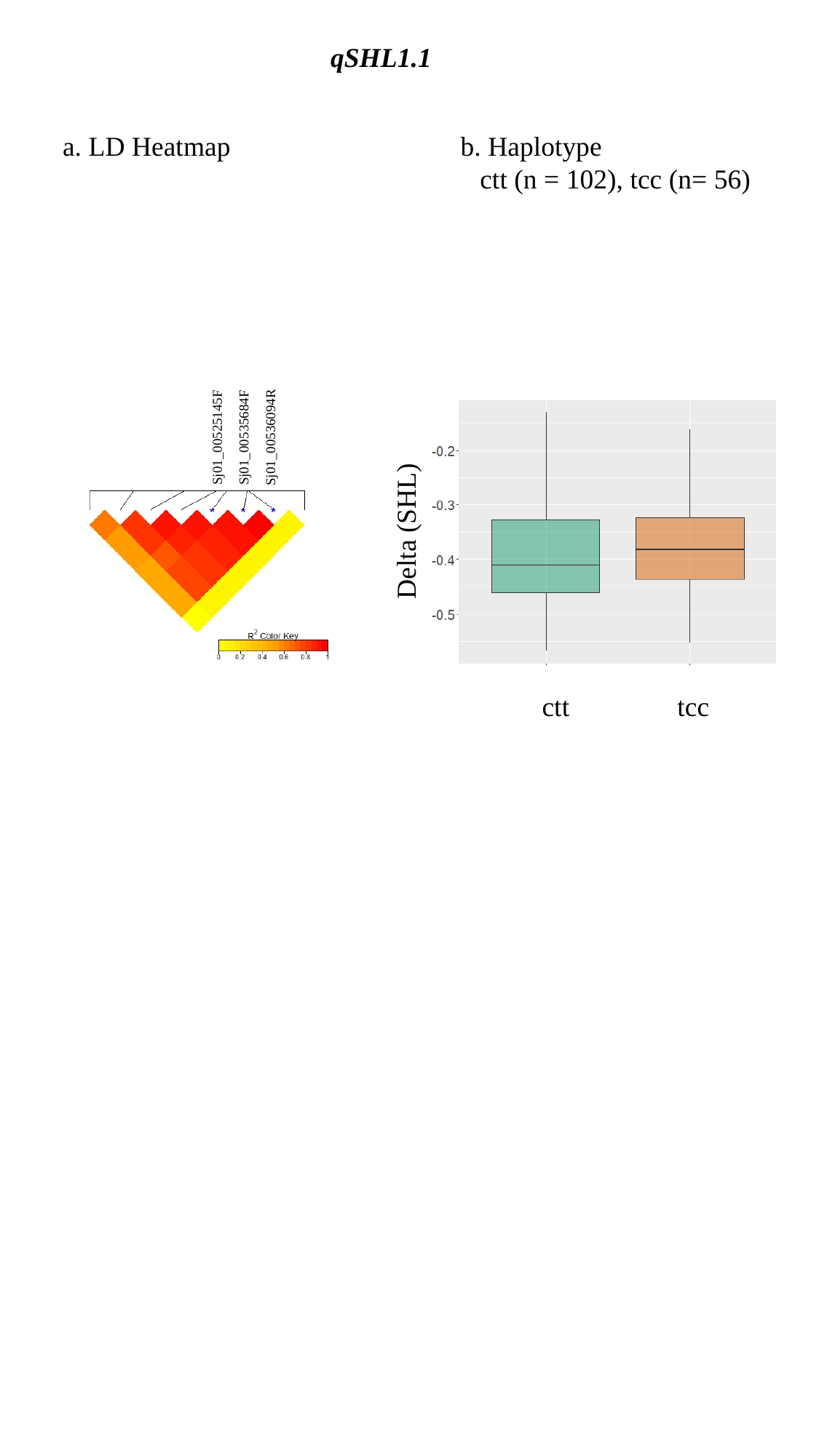

qSHL1.1
a. LD Heatmap
b. Haplotype
ctt (n = 102), tcc (n= 56)
Sj01_00525145F
Sj01_00535684F
Sj01_00536094R
Delta (SHL)
ctt
tcc

### Slide 2
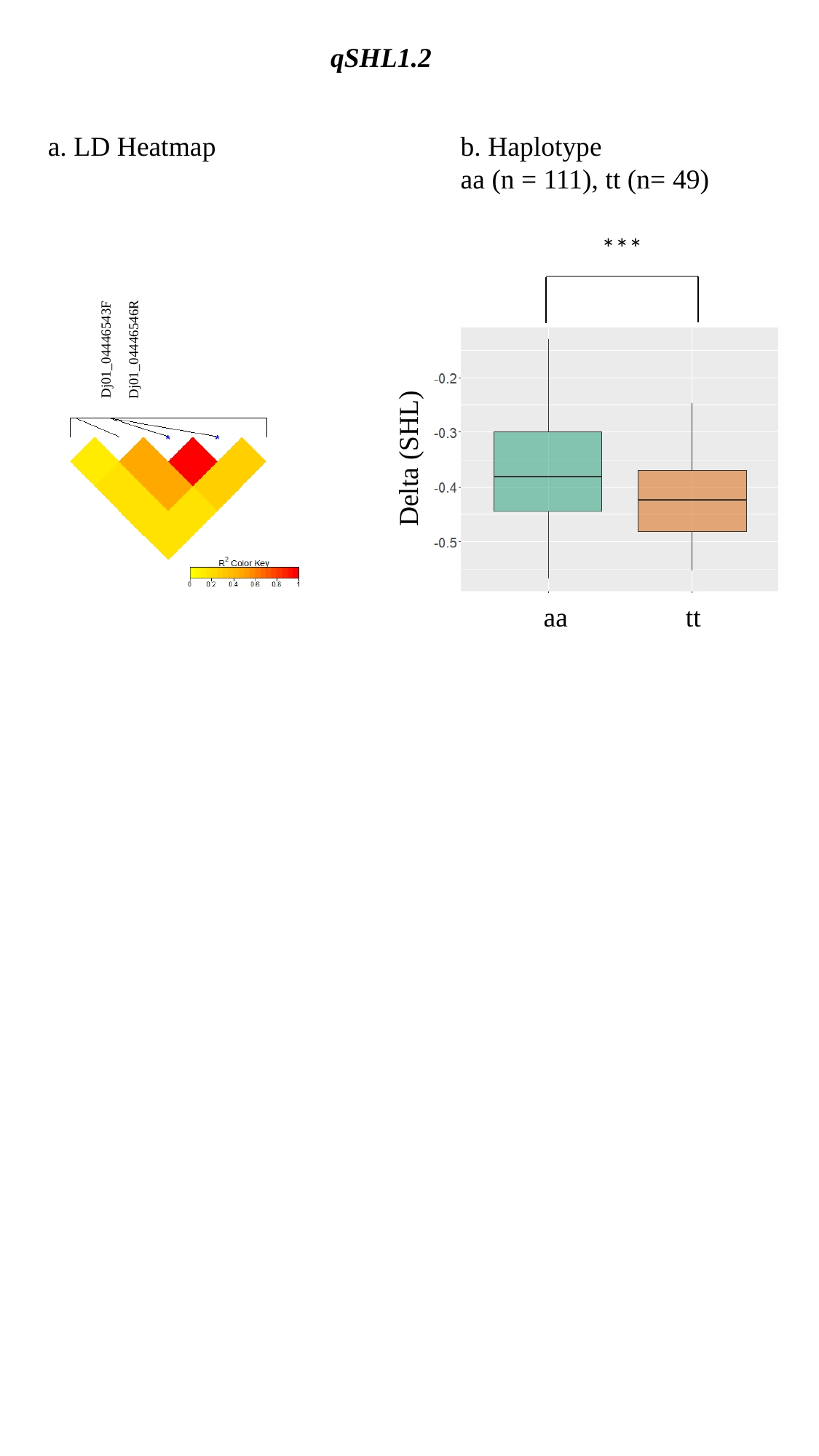

qSHL1.2
a. LD Heatmap
b. Haplotype
aa (n = 111), tt (n= 49)
***
Dj01_04446546R
Dj01_04446543F
Delta (SHL)
aa
tt

### Slide 3
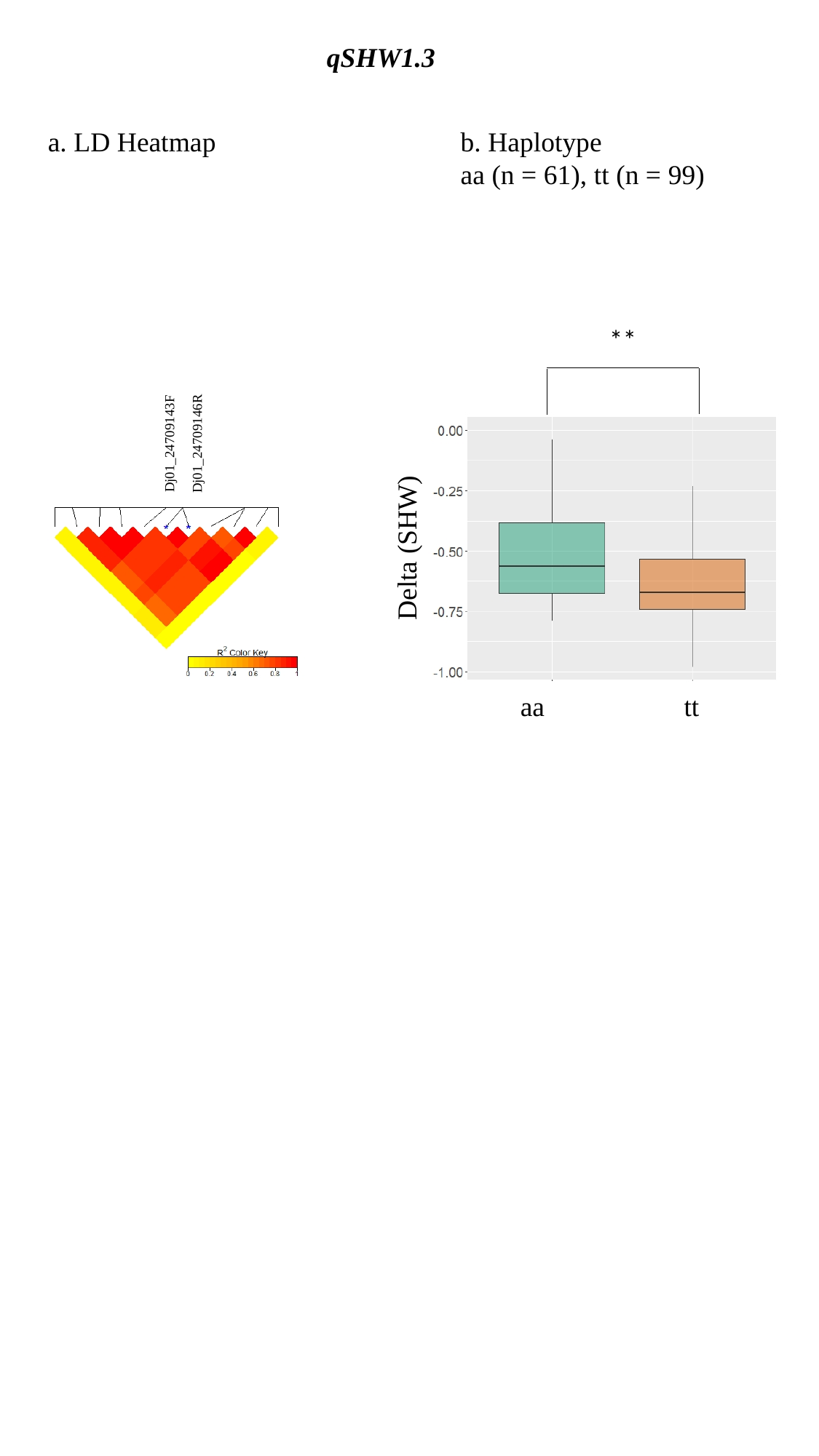

qSHW1.3
a. LD Heatmap
b. Haplotype
aa (n = 61), tt (n = 99)
**
Dj01_24709146R
Dj01_24709143F
Delta (SHW)
aa
tt

### Slide 4
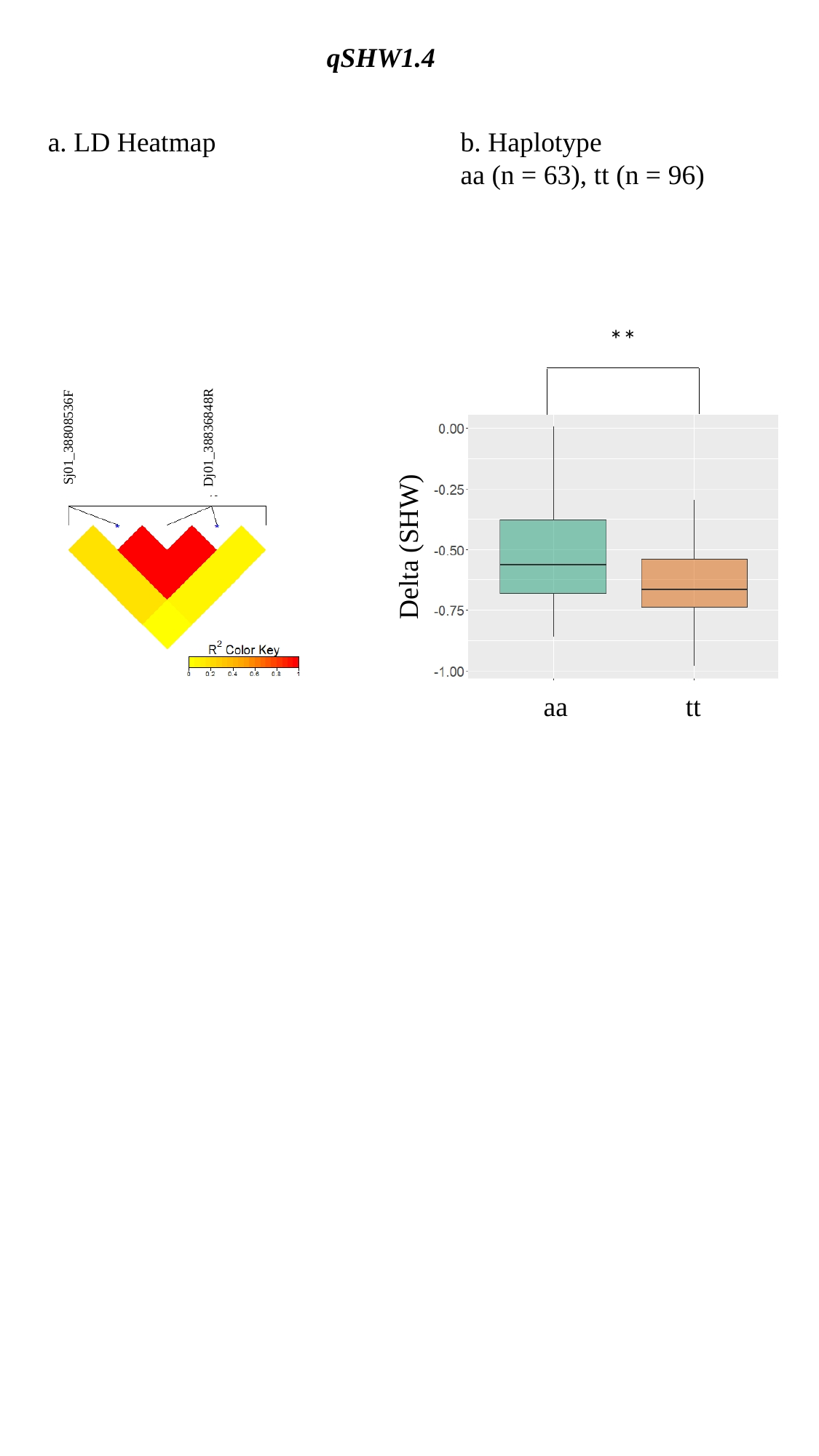

qSHW1.4
a. LD Heatmap
b. Haplotype
aa (n = 63), tt (n = 96)
**
Sj01_38808536F
Dj01_38836848R
Delta (SHW)
aa
tt

### Slide 5
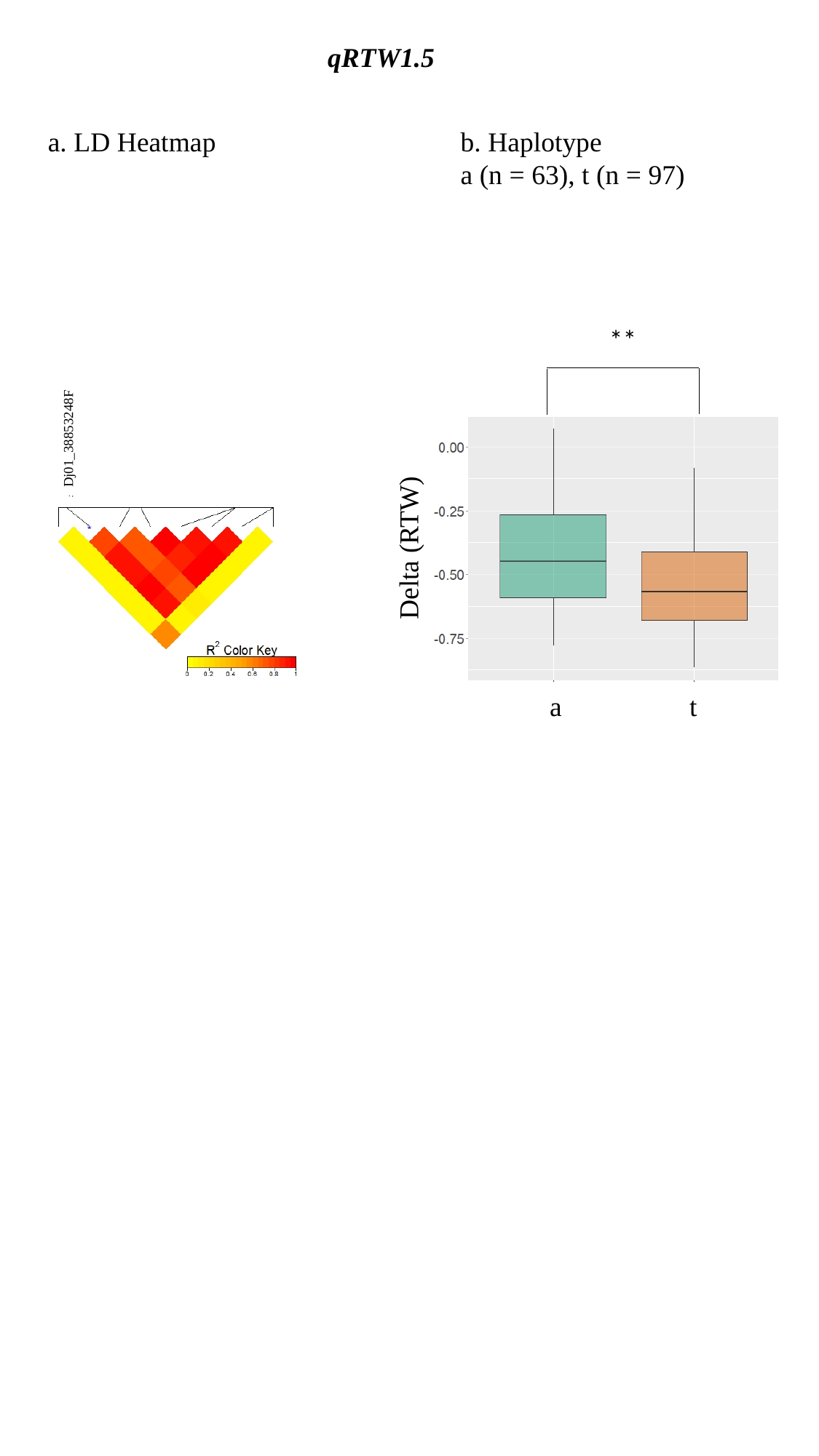

qRTW1.5
a. LD Heatmap
b. Haplotype
a (n = 63), t (n = 97)
**
Dj01_38853248F
Delta (RTW)
a
t

### Slide 6
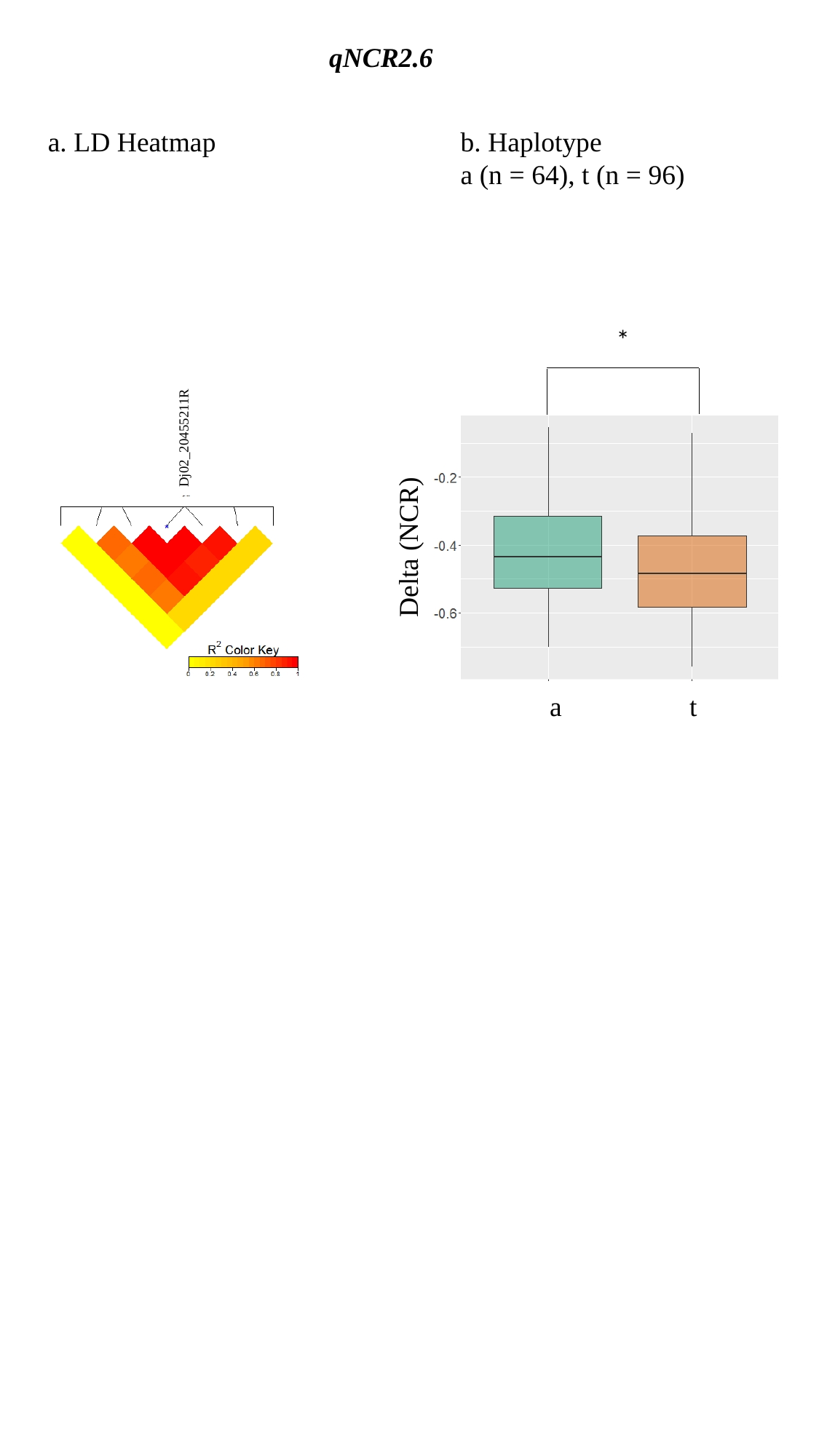

qNCR2.6
a. LD Heatmap
b. Haplotype
a (n = 64), t (n = 96)
*
Dj02_20455211R
Delta (NCR)
a
t

### Slide 7
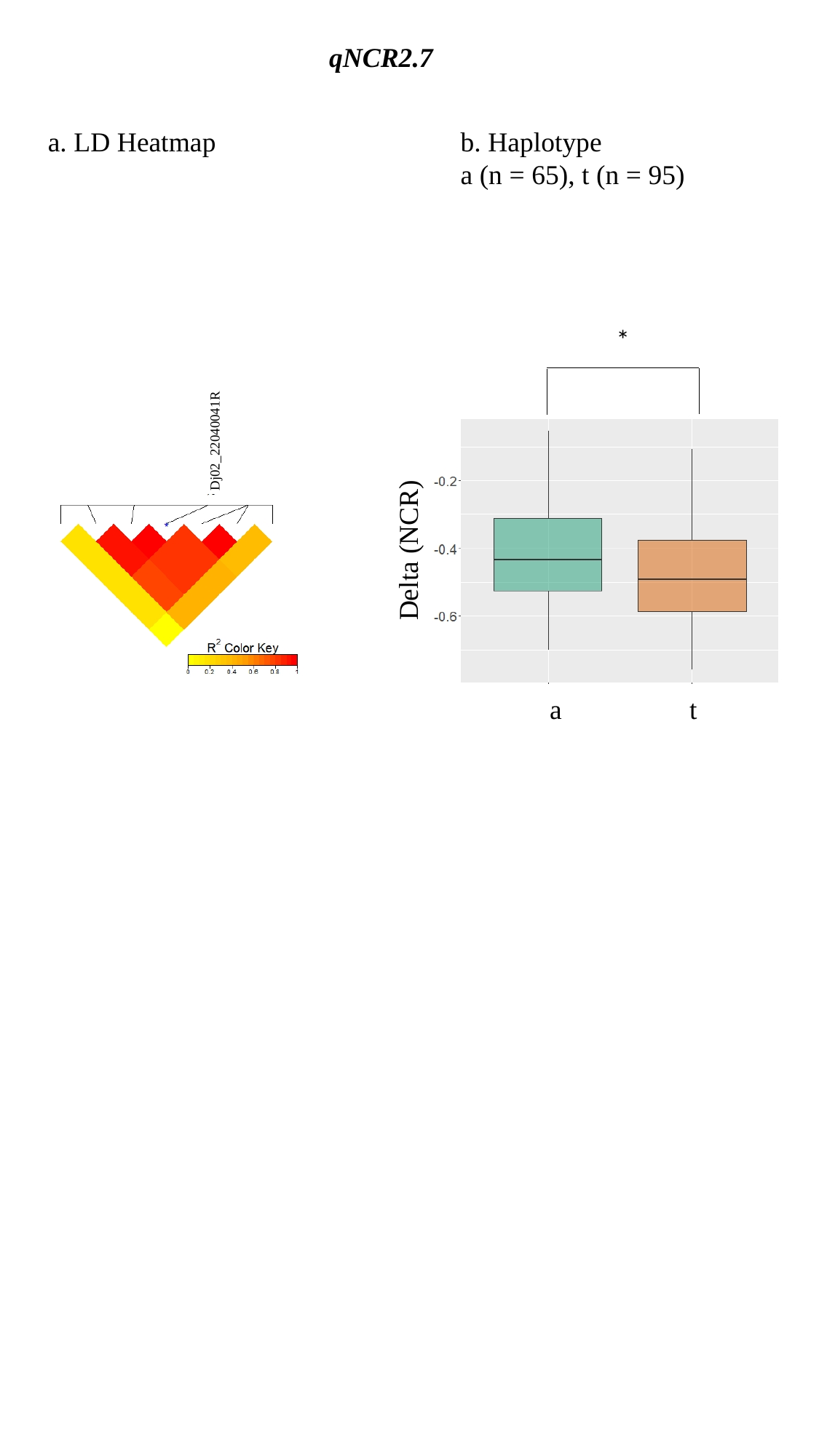

qNCR2.7
a. LD Heatmap
b. Haplotype
a (n = 65), t (n = 95)
*
Dj02_22040041R
Delta (NCR)
a
t

### Slide 8
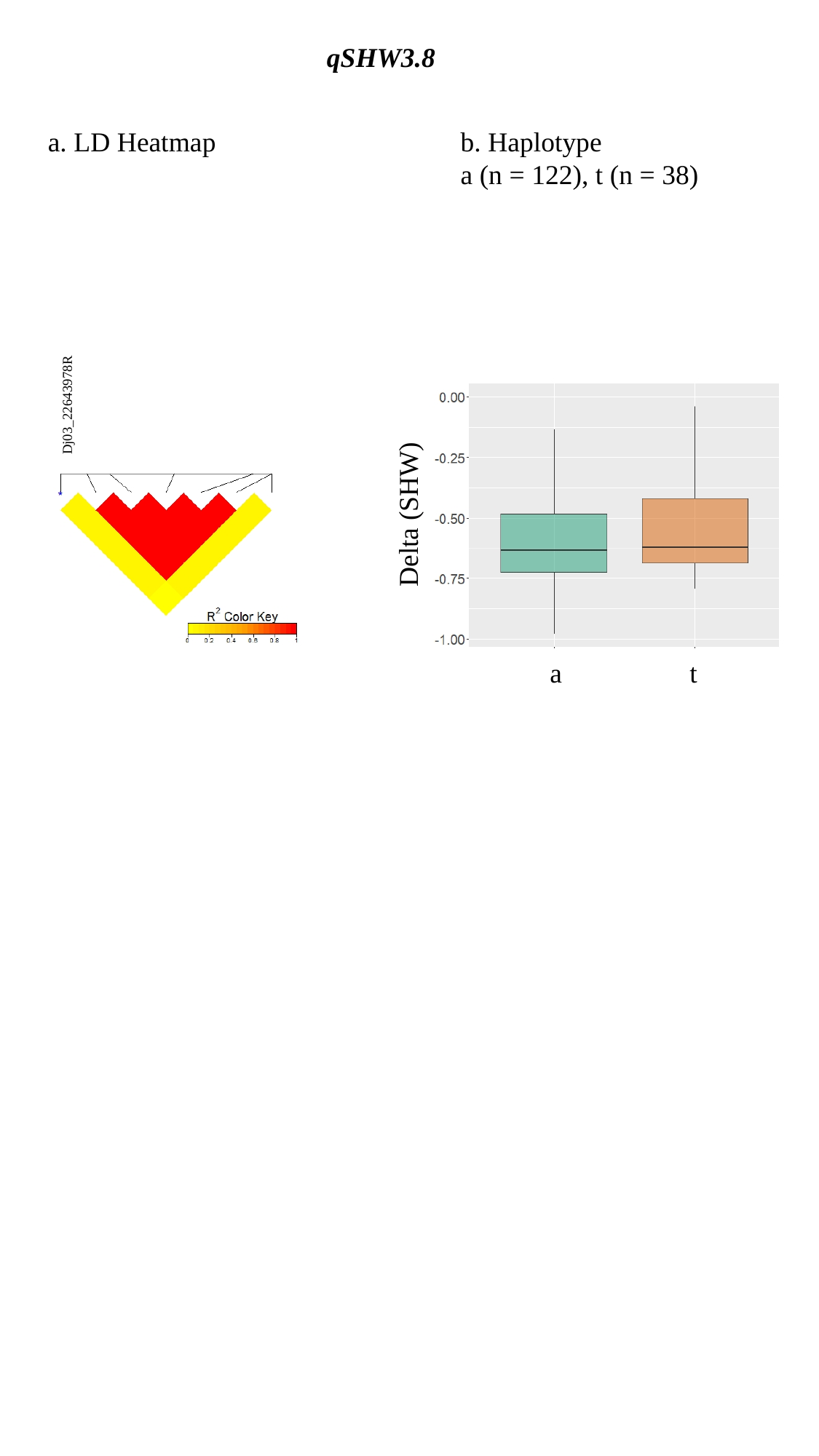

qSHW3.8
a. LD Heatmap
b. Haplotype
a (n = 122), t (n = 38)
Dj03_22643978R
Delta (SHW)
a
t

### Slide 9
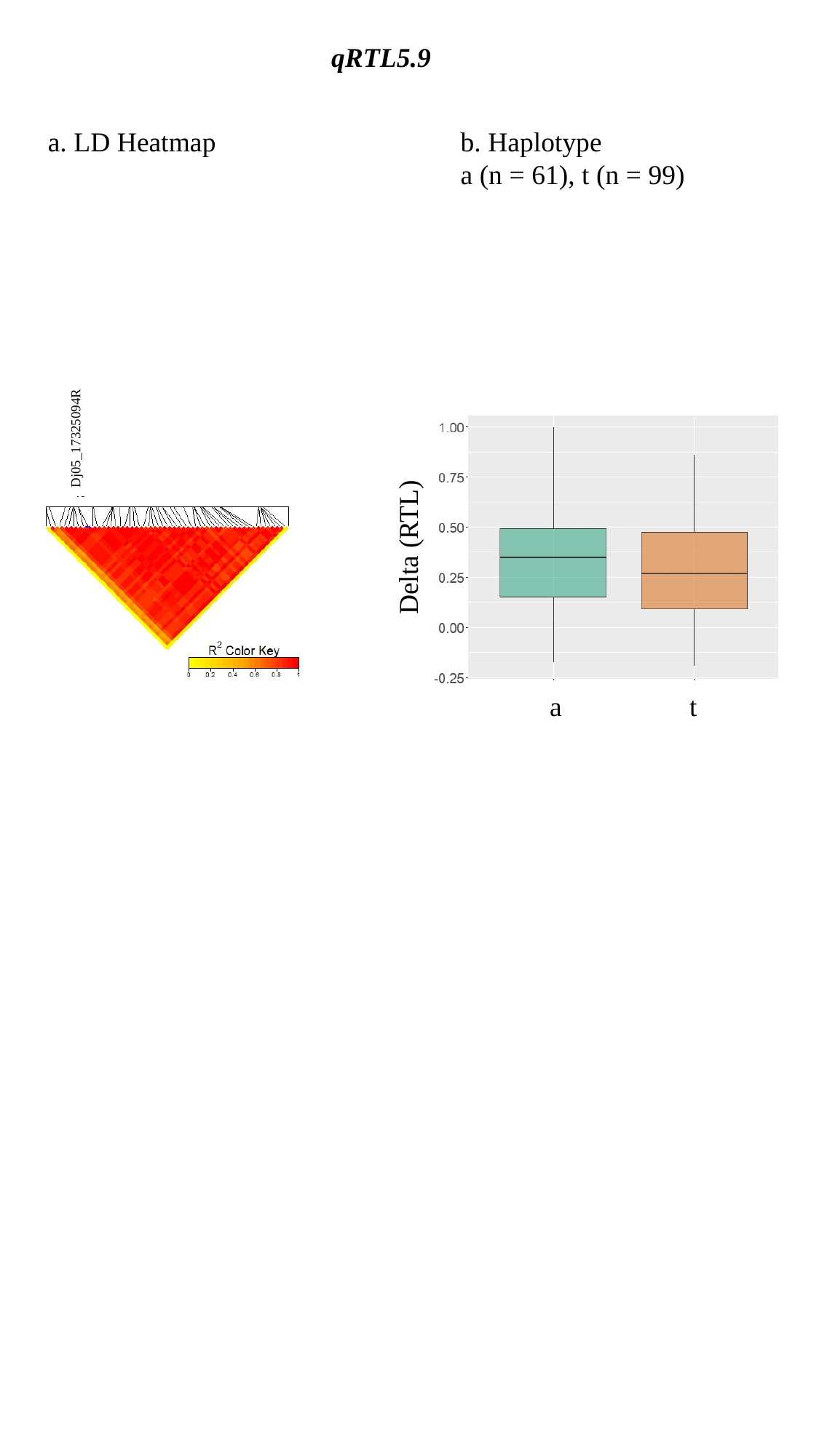

qRTL5.9
a. LD Heatmap
b. Haplotype
a (n = 61), t (n = 99)
Dj05_17325094R
Delta (RTL)
a
t

### Slide 10
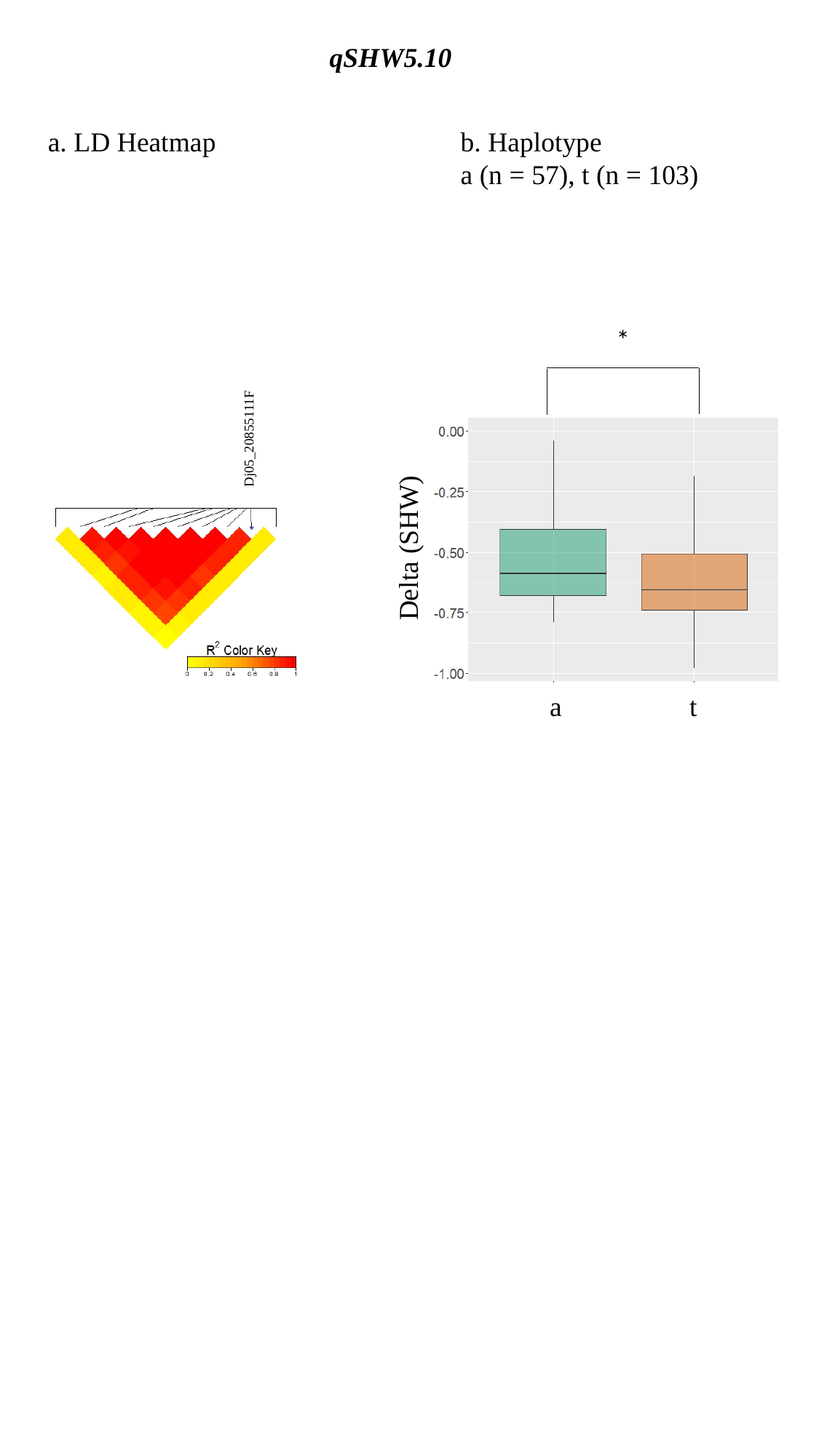

qSHW5.10
a. LD Heatmap
b. Haplotype
a (n = 57), t (n = 103)
*
Dj05_20855111F
Delta (SHW)
a
t

### Slide 11
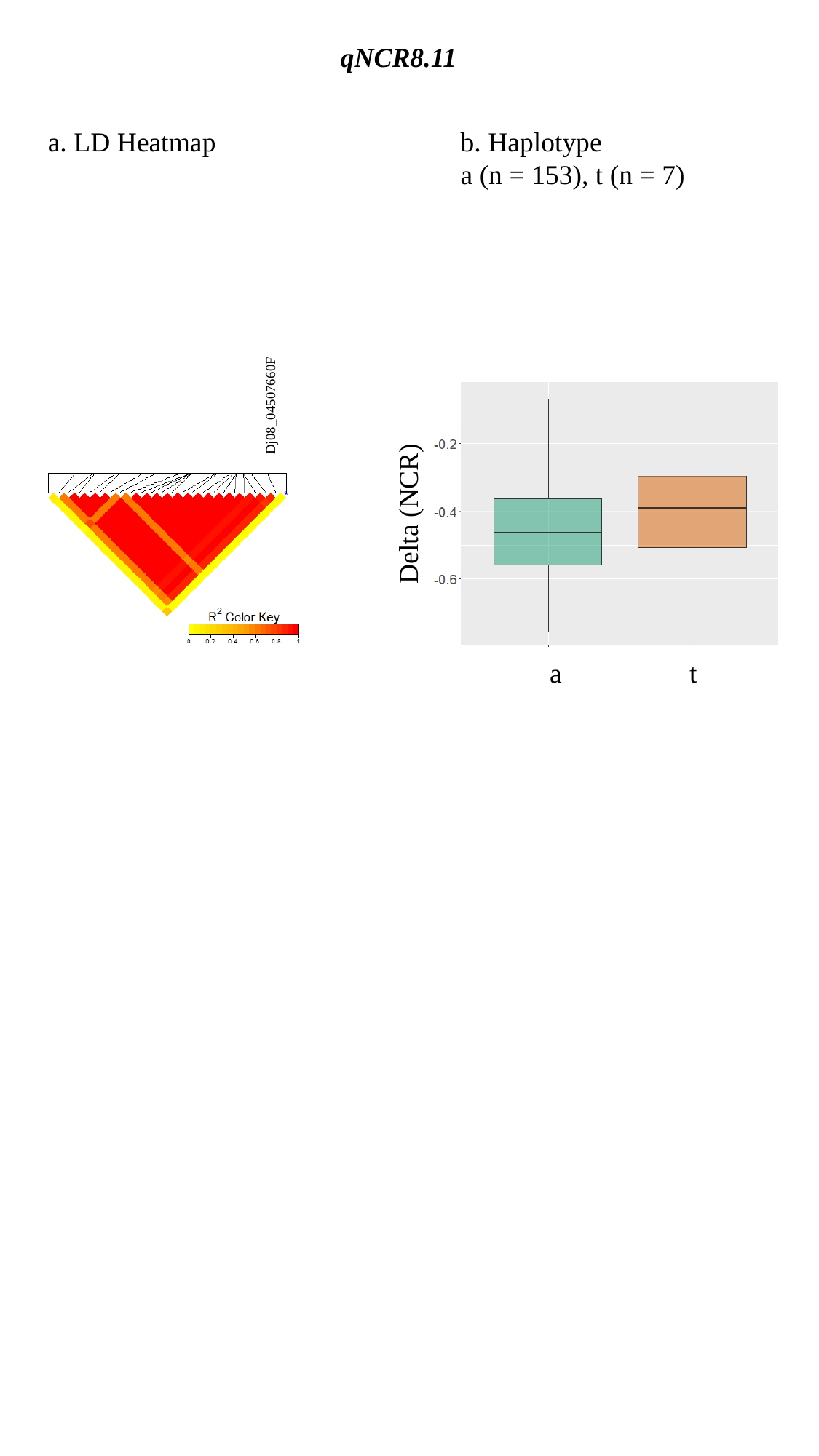

qNCR8.11
a. LD Heatmap
b. Haplotype
a (n = 153), t (n = 7)
Dj08_04507660F
Delta (NCR)
a
t

### Slide 12
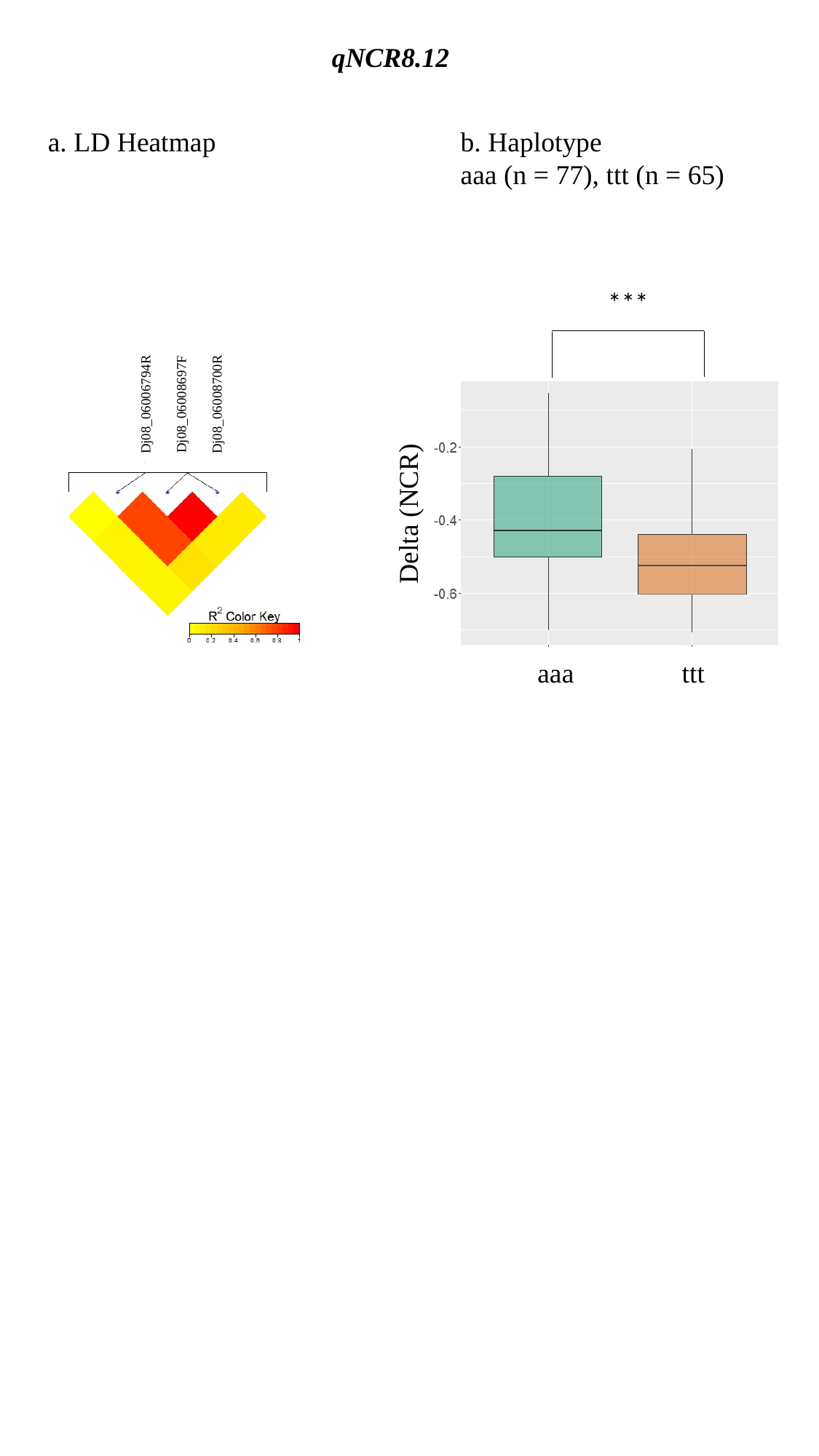

qNCR8.12
a. LD Heatmap
b. Haplotype
aaa (n = 77), ttt (n = 65)
***
Dj08_06006794R
Dj08_06008697F
Dj08_06008700R
Delta (NCR)
aaa
ttt

### Slide 13
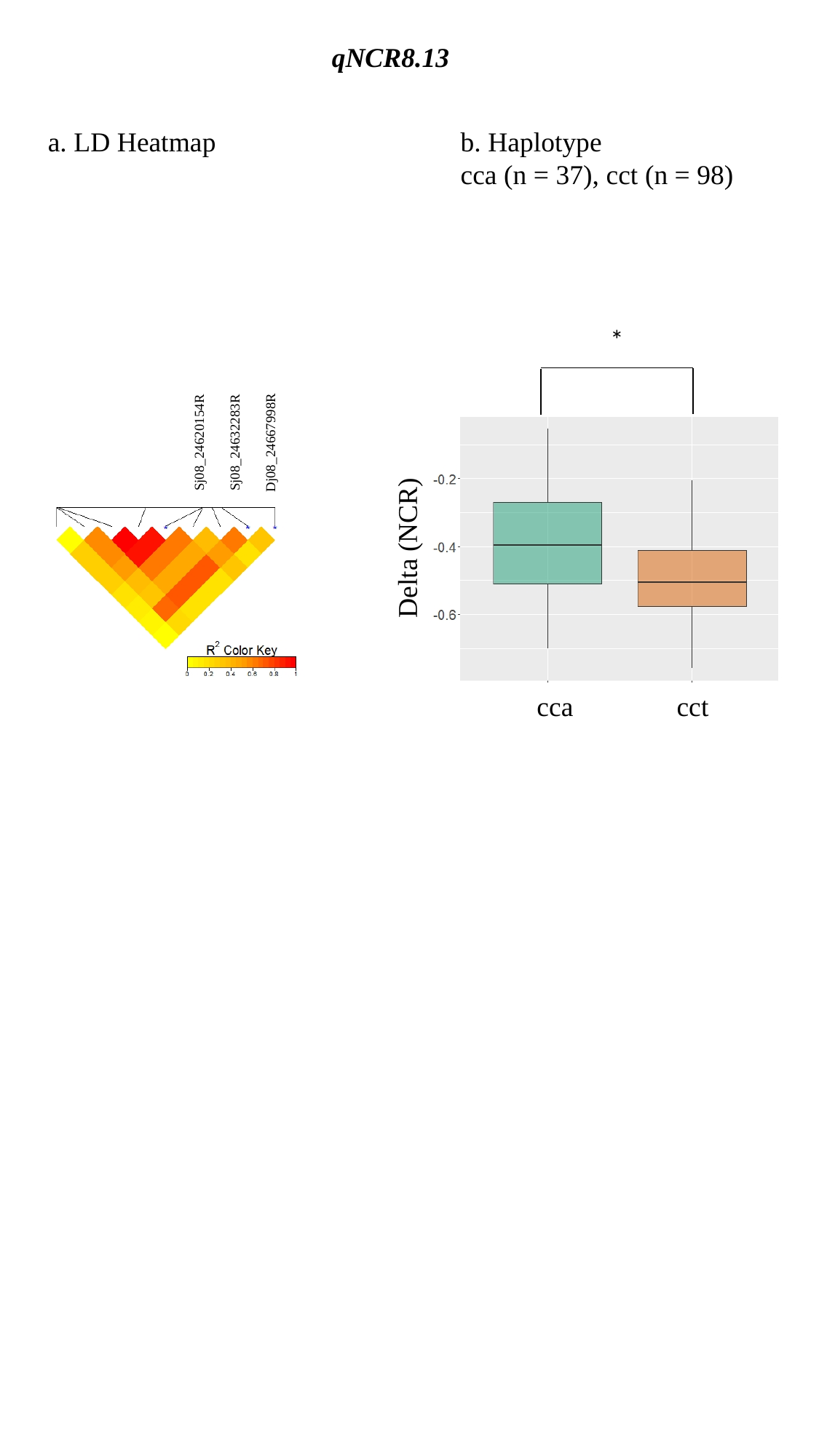

qNCR8.13
a. LD Heatmap
b. Haplotype
cca (n = 37), cct (n = 98)
*
Sj08_24620154R
Sj08_24632283R
Dj08_24667998R
Delta (NCR)
cca
cct

### Slide 14
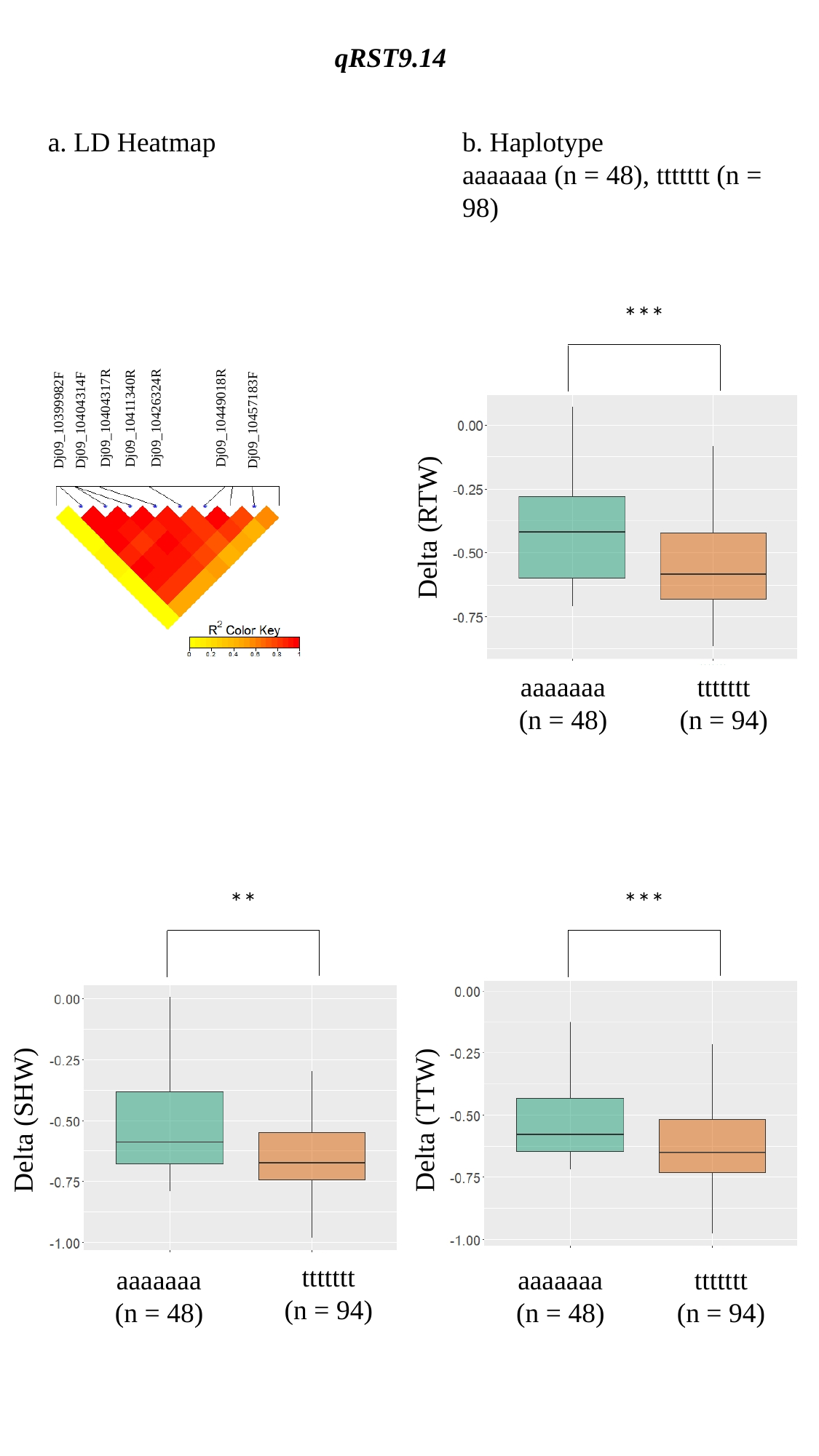

qRST9.14
a. LD Heatmap
b. Haplotype
aaaaaaa (n = 48), ttttttt (n = 98)
***
Dj09_10449018R
Dj09_10404317R
Dj09_10411340R
Dj09_10426324R
Dj09_10404314F
Dj09_10457183F
Dj09_10399982F
Delta (RTW)
aaaaaaa
(n = 48)
ttttttt
(n = 94)
**
***
Delta (SHW)
Delta (TTW)
ttttttt
(n = 94)
aaaaaaa
(n = 48)
aaaaaaa
(n = 48)
ttttttt
(n = 94)

### Slide 15
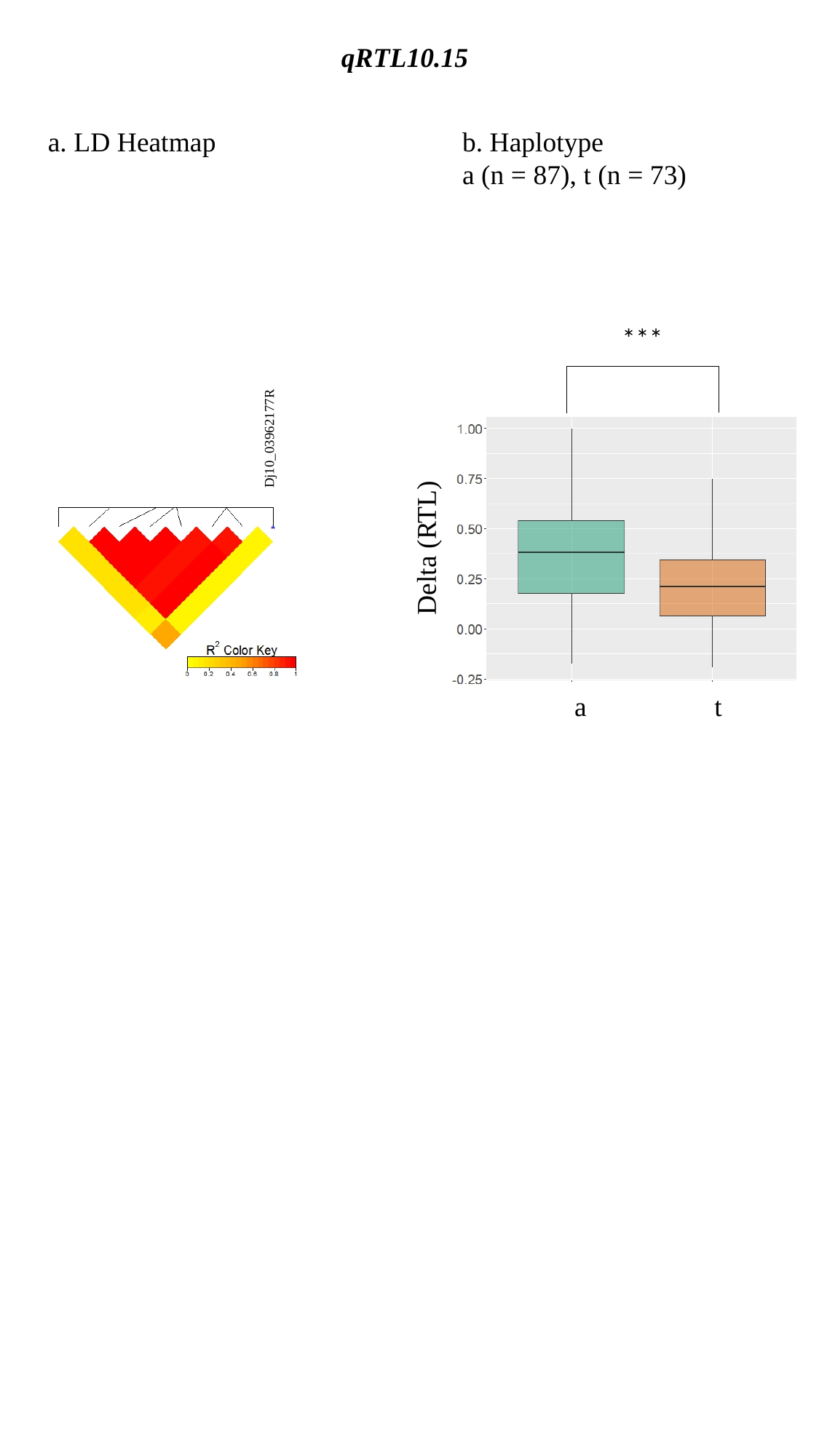

qRTL10.15
a. LD Heatmap
b. Haplotype
a (n = 87), t (n = 73)
***
Dj10_03962177R
Delta (RTL)
a
t

### Slide 16
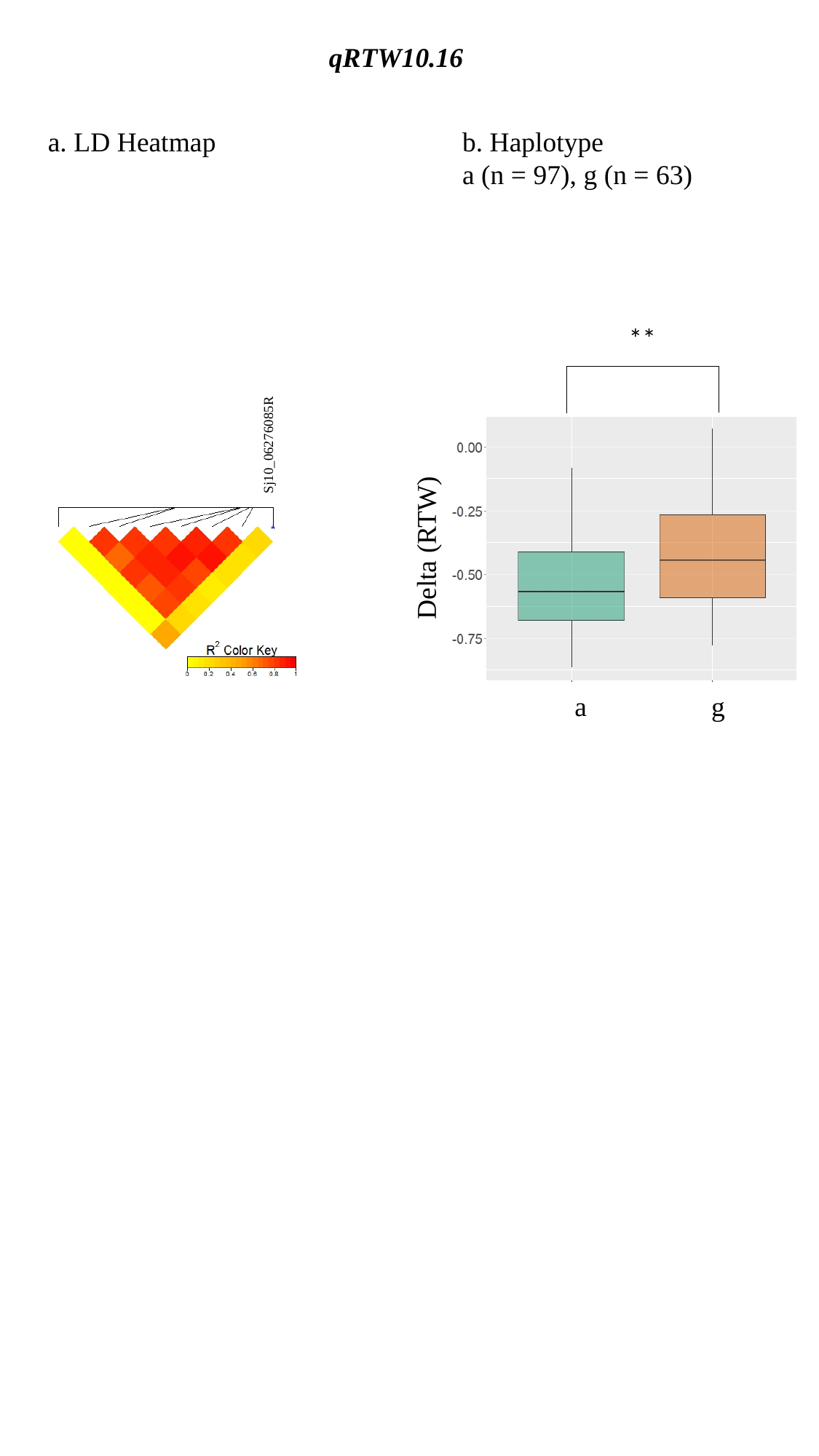

qRTW10.16
a. LD Heatmap
b. Haplotype
a (n = 97), g (n = 63)
**
Sj10_06276085R
Delta (RTW)
a
g

### Slide 17
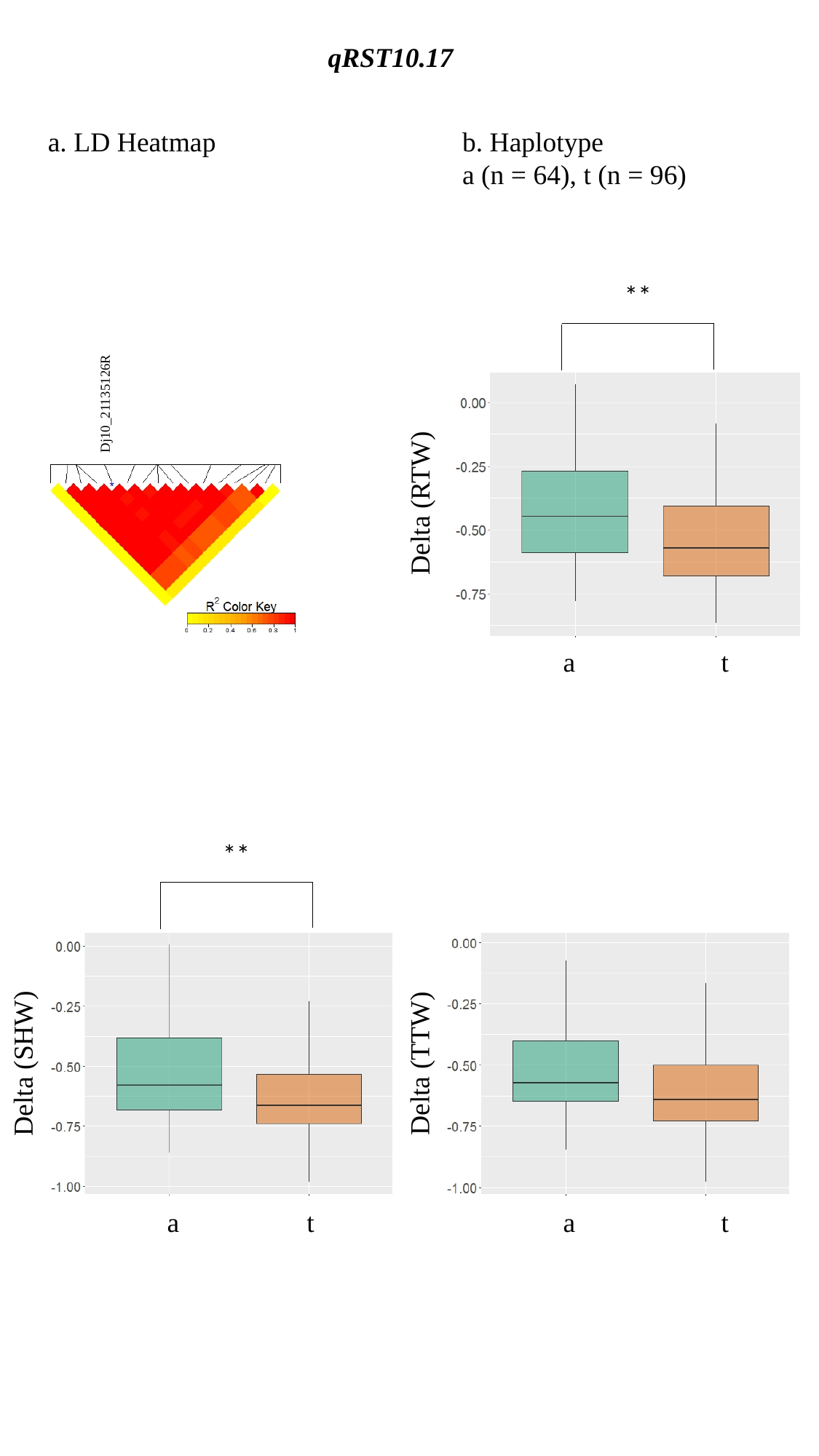

qRST10.17
a. LD Heatmap
b. Haplotype
a (n = 64), t (n = 96)
**
Dj10_21135126R
Delta (RTW)
a
t
**
Delta (SHW)
Delta (TTW)
a
t
a
t

### Slide 18
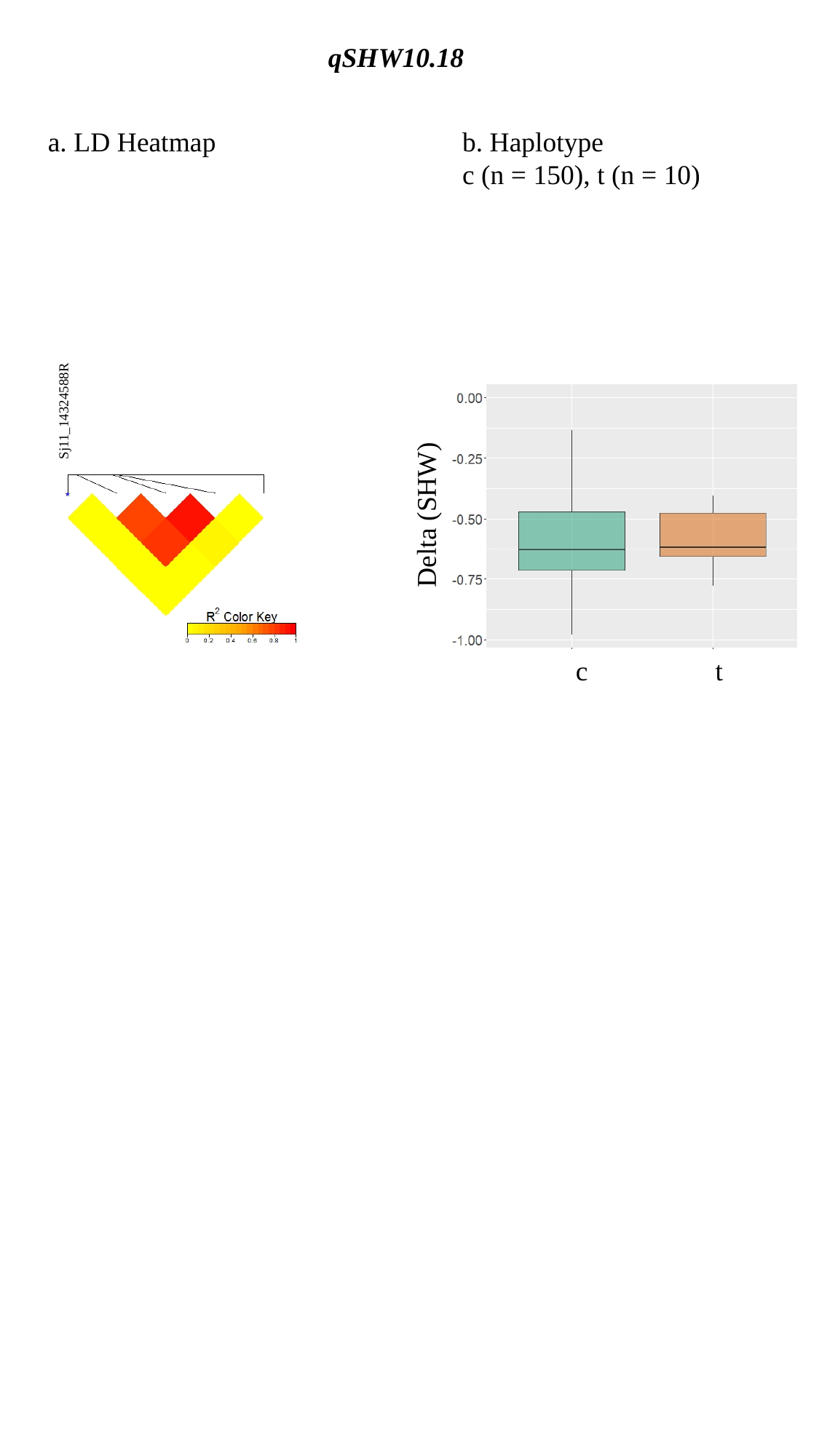

qSHW10.18
a. LD Heatmap
b. Haplotype
c (n = 150), t (n = 10)
Sj11_14324588R
Delta (SHW)
c
t

### Slide 19
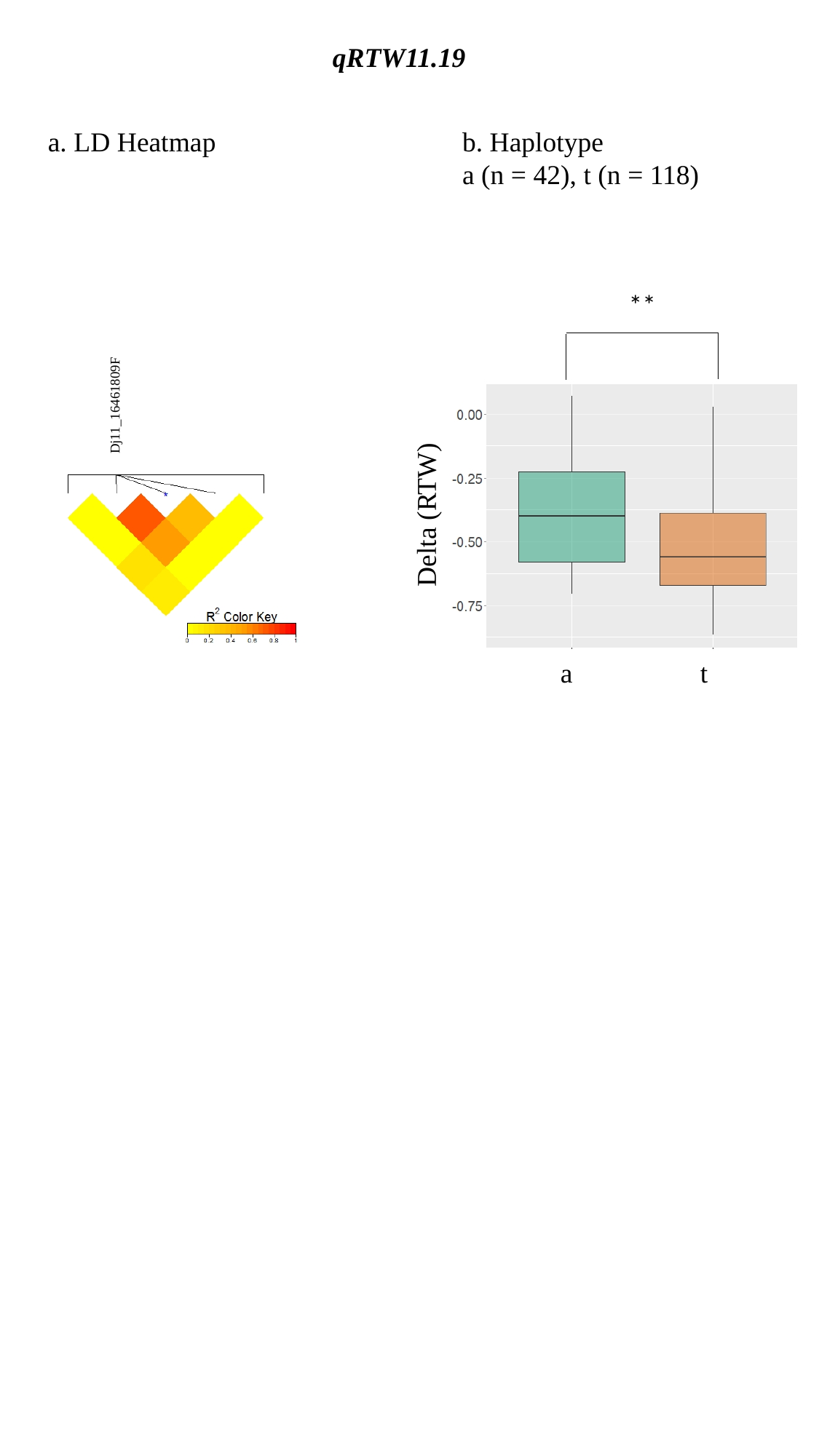

qRTW11.19
a. LD Heatmap
b. Haplotype
a (n = 42), t (n = 118)
**
Dj11_16461809F
Delta (RTW)
a
t

### Slide 20
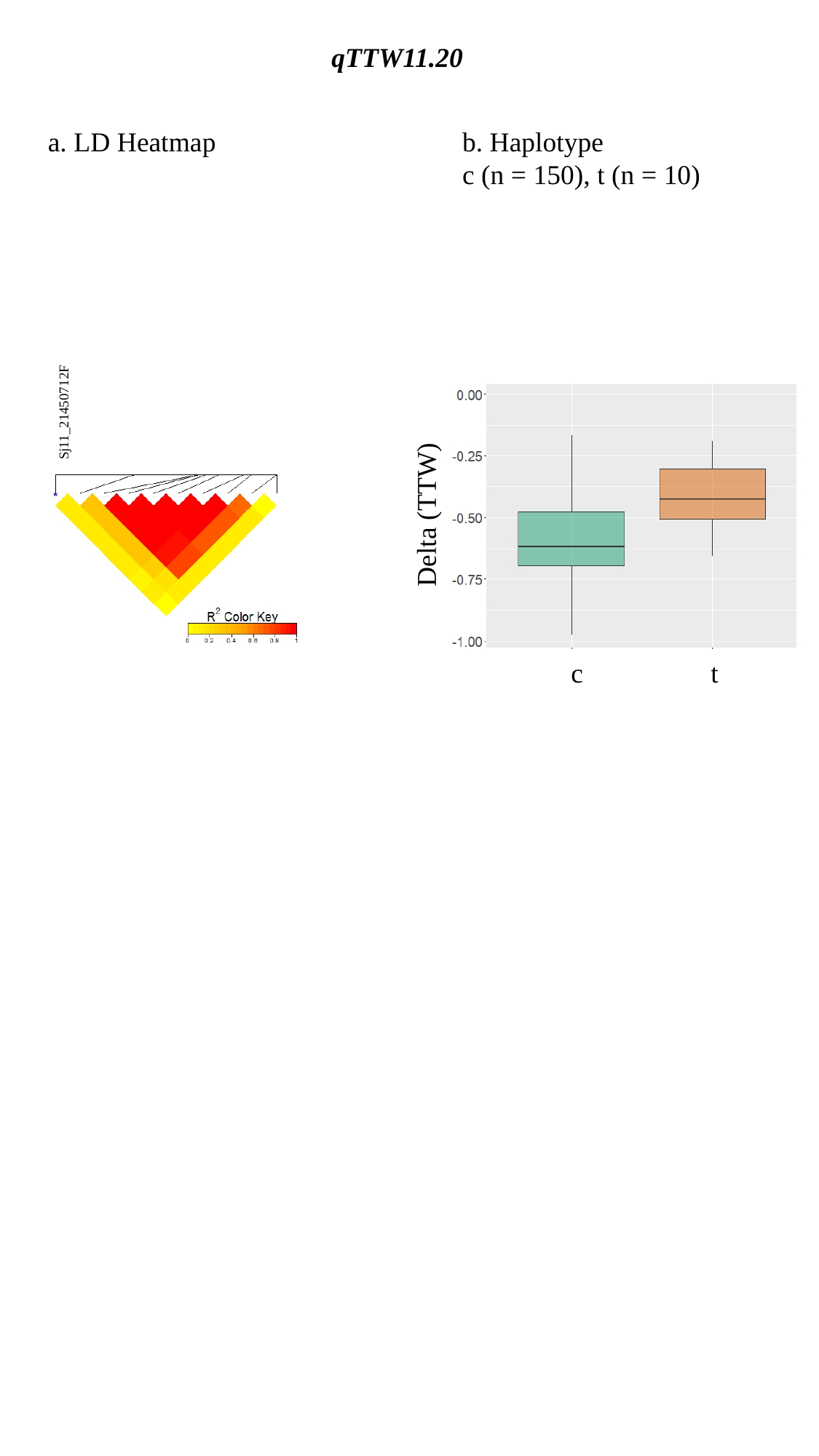

qTTW11.20
a. LD Heatmap
b. Haplotype
c (n = 150), t (n = 10)
Sj11_21450712F
Delta (TTW)
c
t

### Slide 21
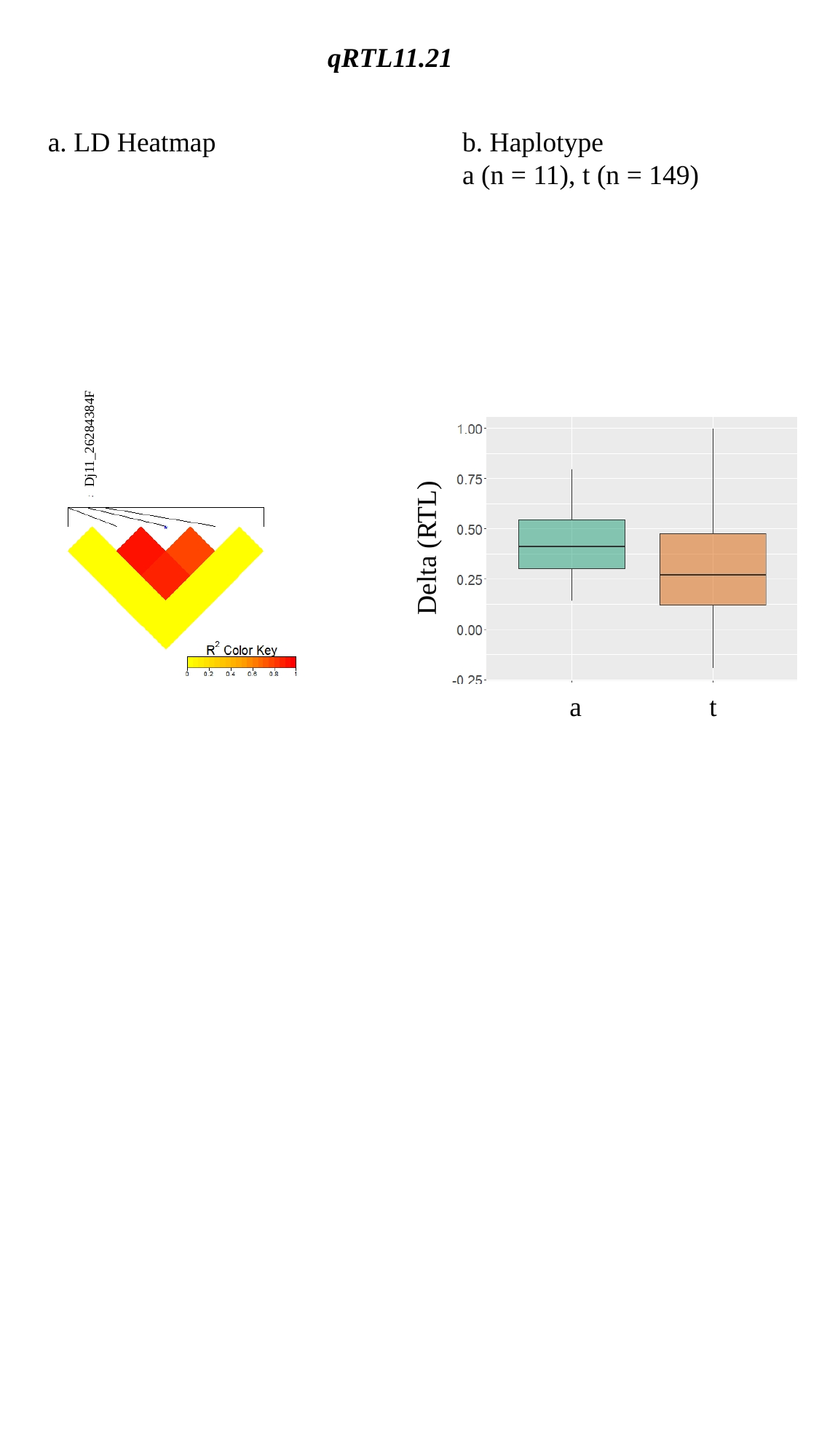

qRTL11.21
a. LD Heatmap
b. Haplotype
a (n = 11), t (n = 149)
Dj11_26284384F
Delta (RTL)
a
t
